# The “Plural Values of Nature” scale: An integrated psychometric scale to measure instrumental, intrinsic and relational values

**DOI:** 10.1101/2025.01.10.632251

**Authors:** Arjen E. Buijs, Sabrina Dressel, Jette B. Jacobsen, Tim M. de Kruiff, Laura Quintero Uribe, Angus Monro Smith, Christian Gamborg

## Abstract

Sustainability transformations critically depend on understanding, assessing and monitoring the plural values of nature held across societies. However, while conceptual frameworks increasingly acknowledge a plurality of values, empirical indicators to reliably assess these multiple value types and their sub-dimensions remain scarce. We introduce the Plural Values of Nature (PVN) scale, integrating intrinsic, instrumental, and relational values into one multi-dimensional scale. This psychometric scale consists of 11 items and is designed to capture both distinctiveness and overlap of values. Using a seven-step process that includes qualitative and cognitive interviews, expert evaluations, and quantitative pre-testing and validation the scale is refined and validated across five European countries (representative sample; *n* = 1028). The validation study shows moderate to high internal consistency, reliability, and validity of the scale across all countries. Based on the development and validation of the PVN, we discuss methodological and conceptual issues and unpack the plurality of values of nature. Initial results show that considering the plurality of values is important to understand people’s attitudes towards e.g., nature restoration: People that endorse multiple values show more positive attitudes towards nature restoration than people endorsing only one specific value type (either instrumental, intrinsic, or relational values), with people endorsing all three value types simultaneously showing the highest support for such policies. As a concise survey-based social indicator, the PVN scale can be used to take stock of the plural values that societies hold regarding the future development of nature and to monitor potential shifts over time.

## 1. Introduction

### 1.1 Background

The world is facing unprecedented environmental challenges, especially related to climate change and biodiversity loss. The Kunming-Montréal Global Biodiversity Framework (GBF) in 2022 has established biodiversity safeguarding targets to be reached in 2030 as well as targets to ensure sustainable use of nature (Hughes & Grumbine, 2023). In doing so, the GBF recognizes that human societies and nature are intertwined, an acknowledgement that is also made in the scientific literature concerning social-ecological systems (Li et al., 2025). To measure development as well as possible future transformation paths, it is important to have indicators that can measure natural as well as societal aspects. Recent syntheses of the diverse values literature argue that value indicators should not be limited to biophysical or monetary measures, but should also include socio-cultural indicators that capture people–nature relationships and the plurality of ways in which nature matters to people (Pascual et al., 2023). However, current biodiversity monitoring and policy practice still rely predominantly on biophysical and, to a lesser extent, monetary indicators. This highlights the need for reliable social indicators that can assess and monitor plural values of nature across populations and over time. Against this background, this paper addresses the societal side of sustainability indicators by proposing and testing the PVN scale as a survey-based social indicator of plural values of nature based on recent theoretical developments in the plural values literature. While some studies (e.g. Wang et al., 2025) build on the ecosystem service framework, or well-being in a broader sense (Li et al., 2025) we go a bit broader, to capture the pluralism of values.

A societal transformation can, in a reasonably well-functioning democracy, only take place if it has public support (Buijs et al., 2025). This support, in part, depends on people’s perceptions, values and cognitions (Patterson et al., 2017). However, views on the value of nature vary within and between sociodemographic groups, communities, cultures, countries and regions (Buijs et al., 2009; Kloek et al., 2017). The IPBES Values Assessment (IPBES, 2022) and the IPBES Transformative Change Assessment (IPBES, 2024) argue for the need to explicitly acknowledge and explore the many different values of nature. Incorporating a wide spectrum of values and releasing latent “nature-positive values” is suggested as an important leverage point for sustainable transformations (Horcea-Milcu et al., 2023; IPBES, 2024). Recognition of the plural values of nature allows for the inclusion of diverse groups and worldviews in environmental management (Chan et al., 2016) and addresses the widely acknowledged need to increase sensitivity to justice impacts of sustainable transformations (Bennett et al., 2019). Potential scenarios for transformative futures have also increasingly engaged with the plural values of nature. For example, the Nature-Futures Framework (NFF) has been developed as a tool to explore integrative paths for bringing about transformative change, focusing on positive and diverse relationships between people and nature (Pereira et al., 2020). The three paths developed in this approach (Nature for People, Nature for Nature, and Nature as Culture) are explicitly inspired by the plurality of values of nature and are increasingly used in scenario development, assessment and implementation focused on inclusive transformative change, sensitive to the diverse values of nature endorsed by stakeholders and local communities (e.g. Durán et al., 2023; Lengieza et al., 2023; Quintero-Uribe et al., 2022). This increased focus on the relevance of the plural values of nature, and the extensive theoretical and conceptual developments over the last years in understanding these concepts (IPBES, 2022), suggests the need for understanding how the plurality of values can be measured and incorporated in inclusive environmental policies. In this paper we engage with this need through developing and testing a psychometric scale that empirically measures the plural values of nature and can function as a survey-based social indicator for assessing and monitoring these values across populations.

### 1.2 Values of Nature

To understand the richness of people’s relationships to non-human nature, the IPBES Value assessment argues for the need to distinguish worldviews, knowledge systems, broad values and specific values. Where broad values are generally considered to be stable, specific values are more changeable over time (IPBES, 2022). In this paper, we focus on the specific values of nature (Ibid.).

Specific values of nature have been conceptualised into three broad categories - instrumental, intrinsic and relational values (Chan et al., 2016; IPBES, 2022; Muraca, 2011; Pascual et al., 2023). However, decades of philosophical, sociological and psychological explorations of their ontological, epistemological, conceptual, and empirical underpinnings (e.g. Buijs, 2009; Chan et al., 2018; R. Gould et al., 2024; Himes et al., 2024; Kempton et al., 1995; Lockwood, 1999; Muraca, 2011; Stålhammar & Thorén, 2019) have shown that this categorisation is not without issues. In particular, for psychometric scale development it is important to acknowledge that the three categories are not easily distinguishable or separated. As highlighted in recent scholarship (Himes et al., 2024; Kenter et al., 2019), fuzzy boundaries exist between them, with a high degree of overlap. Moreover, many individuals simultaneously endorse multiple values from each of the categories, as shown in the applied literature (e.g. Uggeldahl et al, 2025).

Instrumental values can be conceptualised as being based on the value that an individual and/or society may derive *from* nature, that is, from animals, plants, ecosystems and natural processes (Himes & Muraca, 2018). Instrumental values are the values of living entities as means to achieve human-defined ends, or to *satisfy human preferences* (Pascual et al., 2017), or the basis of *services* and *benefits* that nature can provide for human beings (See, 2020). Especially the latter is often associated with the concept of ecosystem services, providing regulating, cultural, and supporting services (Feucht et al., 2023). One important characteristic of instrumental values is that they are transactional, and in principle substitutable, though not always in practice (Himes et al., 2024).

Intrinsic values have been explored extensively in environmental philosophy. Central to the varieties of the understanding of the concept are three key elements: i) the *inherent*, non-relational properties of nature and/or natural objects; ii) the value of nature *independent* of human valuation and judgment, and iii) natural entities as bearer of inherent moral value as ends in themselves, and subjects/objects with their own good (O’Neill, 1992). In the IPBES approach, intrinsic value is defined as the “…inherent value, that is the value something has independent of any human experience or evaluation. Such a value is viewed as an inherent property of the entity (e.g., an organism) and not ascribed or generated by external valuing agents (such as human beings).” (Pascual et al., 2017: 14). Unlike instrumental values, intrinsic values cannot be substituted.

Relational values as a concept has emerged in the last decade in research and practice related to nature and biodiversity (Gould et al., 2024), as well as in policy. Relational values are values of nature stemming from, often reciprocal, relationships (interactions, responsibilities) between people and nature or among people through, or with regards to, nature (Himes et al., 2024). Importantly, relational values have been suggested as a way to break up the dichotomy of instrumental and intrinsic values. The notion was first articulated in land-use literature by Brown (Brown, 1984) as ‘relational realm’ which is called a value concept, though importantly not as a suggested third value type (Chan et al., 2018). As a value type proper, it was coined in the philosophical literature as part of a new axiological framework by Muraca (2011) and in a broader sustainability science context by Diez et al. (2015) and Chan et al. (2016). The advent of this suggested type of value can be seen as a response to a growing critique of (economic) environmental valuation (e.g. Spash, 2020) and as an effort to move beyond the heavily debated human-nature dichotomy (Himes et al., 2024). It also responds to critiques that instrumental and intrinsic values only capture a narrow spectrum of possible values (R. K. Gould et al., 2024; Himes et al., 2024). The concept of relational values is still under debate for various reasons. Some authors argue that relational value is an addition to instrumental and intrinsic value (James, 2022); others that all environmental values are ultimately relational, or at least that relationality is inherent to instrumental and intrinsic values (Luque-Lora, 2023; Norton & Sanbeg, 2021). When the term relational values is used, it aims to capture a more diverse and equitable array of nature’s values than intrinsic and instrumental values - meanings and uses, including e.g., kinship, stewardship, indigenous, spiritual, and place-based values, as well as eudemonic values; i.e., living a fulfilling life of care, well-being and reciprocity (Anderson et al., 2022; Chan et al., 2018; Feucht et al., 2023; Himes et al., 2024; Pratson et al., 2023). An argument for its use is that relational values can be seen as distinct from instrumental and intrinsic values in different ways: relationships have inherent interconnected qualities; they are non-substitutable; and relationships can be reciprocal (instrumental values of nature are transactional).

### 1.3 Developing a scale for measuring plural values of nature

The aim of this paper is to develop a psychometrically validated quantitative scale that can serve as a survey-based social indicator for measuring the plurality of values of nature. Here we focus on the Global North, more specifically Europe as clear spatial boundary in which we seek validity for the scale. Such a scale can be used in quantitative surveys and can contribute to our understanding of not only the diversity of values and how they are related, but also how values impact individual attitudes, motivations and preferences for environmental policies, stewardship or activism (Runhaar et al., 2019; Steg, 2016; Taye et al., 2018). A psychometric scale is a set of standardized, multiple items which are designed to measure a specific latent construct in a reliable and valid way; that is, it consistently produces similar results when used repeatedly under similar conditions and accurately measures the latent construct(s) it is intended to measure. Given the inherent fuzziness, however, we expect that some survey items would have cross-loadings across multiple factors. Nevertheless, rather than being a limitation, such cross-loadings are in fact conceptually coherent with the plural and fuzzy nature of values of nature, (Himes et al., 2024) and the scale we develop in this paper is explicitly designed to capture both distinctiveness and overlap. Nonetheless, there is a fundamental trade-off that has to be considered: Whilst a larger number of items will be required to capture complex concepts, the number of items has to be balanced against the potential use and the need for scale brevity in order to maximize response rate in survey research (Robinson, 2018). While validity was the first criterion, we explicitly aimed for a scale limited in length to enhance its uptake in survey research in environmental sciences.

A valid psychometric scale to measure the plurality of values of nature must respond to several criteria. Based on methodological literature on psychometric scales (G. O. Boateng et al., 2018; Hair et al., 2019; Robinson, 2018), conceptual considerations of the plural nature of values of nature with potential fuzzy boundaries (Himes et al., 2024; Pascual et al., 2023) and pragmatic limitations to the length of a scale to be used in real-life empirical research (Stern et al., 1998), we suggest four distinct criteria: First a psychometric scale by definition must be validated conceptually, empirically, and statistically with representative samples of the general public (G. Boateng et al., 2018). Second, in order to understand the plurality of values, the potential conceptual overlap between values as well as potential empirical co-occurrence of multiple values, the scale should measure all three values simultaneously. Third, as the three values potentially co-occur, and may have fuzzy boundaries, trying to distinguish between them will likely be country and/or culture specific. Therefore, the geographical scope of the scale must be clear, and it is crucial to test validity and reliability in different countries within the geographical scope. Fourth and last, the scale must be applicable in real-life research, most notably surveys to diverse publics. Consequently, the number of items must be limited to allow sufficient space for other concepts to be measures in surveys, such as attitudes, beliefs, behaviours, nature experiences etc. (Stern et al., 1998). Obviously, this last point involves some trade-offs with the other criteria – as will also be clear in the proposed scale and feature as an aspect in the discussion.

Various scales and item-batteries exist to measure values of nature (e.g. Feucht et al., 2023; Klain et al., 2017; Lengieza et al., 2023; Lou et al., 2025; Mrotek et al., 2019; See et al., 2020). Meanwhile, most existing scales tend to focus on, or emphasize, only one value of nature, have not been explicitly validated, are developed and/or used with specific samples - most notably students, are focusing on nature as community, or they are developed and validated in only one country or one cultural context. We acknowledge that several well established and cross-culturally validated scales exist which attend to human-nature relationships, but not explicitly targeted at values of nature, such as the revised New Ecological Paradigm scale (NEP) (Dunlap et al., 2000); connectedness to nature scale (Mayer & Frantz, 2004), and the Environmental Concerns scale (Schultz, 2001).

Based on these reflections, our aim is to develop a solid scale in a rigorous manner, with the sole purpose of measuring the plural values of nature. The scale integrates all three values described above as different dimensions within one psychometric instrument. To maximise usability and to increase the potential for inclusion in larger questionnaires (Stern et al., 1998), we aim for a condensed scale validated across five European countries.

## 2. Materials and Methods

We developed the Plural Values of Nature (PVN) scale, following recommended development and validation practices as outlined in Hair et al. (Hair et al., 2019) and Boateng at al. (G. O. Boateng et al., 2018), via seven steps (Fig 1). Table S1 in the supplementary material provides the formulation of all items at each step and the changes made between them. Figure 3 provides the formulation of the items in the final scale. A notable difference from other attempts to develop a psychometric measurement scale is that we developed PVN simultaneously within multiple (European) countries and related cultural settings, which required additional considerations regarding the scale validation and statistical analysis. Thus, during the development and validation steps, special attention was given to existing guidelines for cross-cultural scale adaptations.

**Fig. 1.**
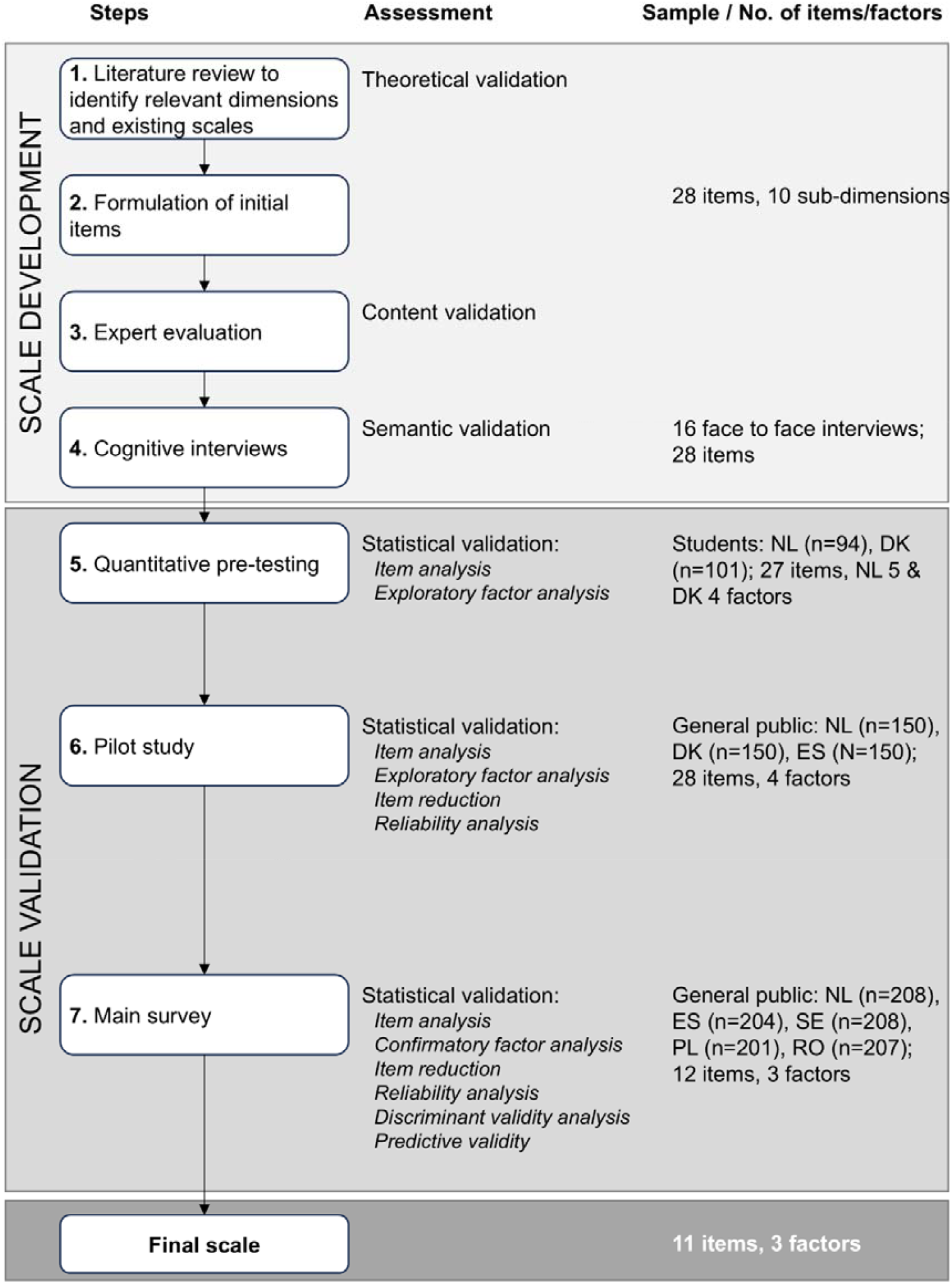
Methodological overview of PVN scale development and validation process. The figure shows the chronological steps taken for scale development and scale validation including assessment procedures, used samples, numbers of items and retained factors.

### 2.1 Instrument Development

An initial narrative literature review was carried out to identify existing related scales and explore relevant subdimensions that should be considered within each value type. For instrumental values, we build on previous conceptualisations of ecosystem services (ESS) and Nature’s Contributions to People (NCP), differentiating between three subdimensions of instrumental values: *material, non-material*, and *regulating contributions* to people (Fig 2)(Diaz et al., 2018). For intrinsic values, we build on existing conceptualisations that distinguish (at least) two subdimensions: *nature as an end in itself* and the recognition that nature has *value beyond human benefit* (Himes et al., 2024). The conceptualisation of relational values proved to be the most difficult, as this relatively new concept is still in development, and is conceptually quite rich, complex, and culturally diverse (R. Gould et al., 2024; Himes et al., 2024). Meanwhile, several item batteries have been developed for use in survey studies (Feucht et al., 2023; Klain et al., 2017; Lengieza et al., 2023), although to the best of our knowledge, only the scale used by Klain et al. (2017) has been used more than once. Next to these existing scales, we found inspiration to distinguish subdimensions for relational values from the IPBES Value Assessment, which differentiates between *place attachment and identity*; *fulfilment and well-being* (eudemonia); *spirituality*; and *care* (as stewardship responsibility) (Anderson et al., 2022). Based on other reviews of relational values (Feucht et al., 2023; Pratson et al., 2023), we added the notion of *kinship* with nonhuman entities as a fifth subdimension. With the three subdimensions for instrumental values and two subdimensions for intrinsic values, this resulted in a structure to develop items consisting of in total ten subdimensions (Fig. 2).

**Fig. 2.**
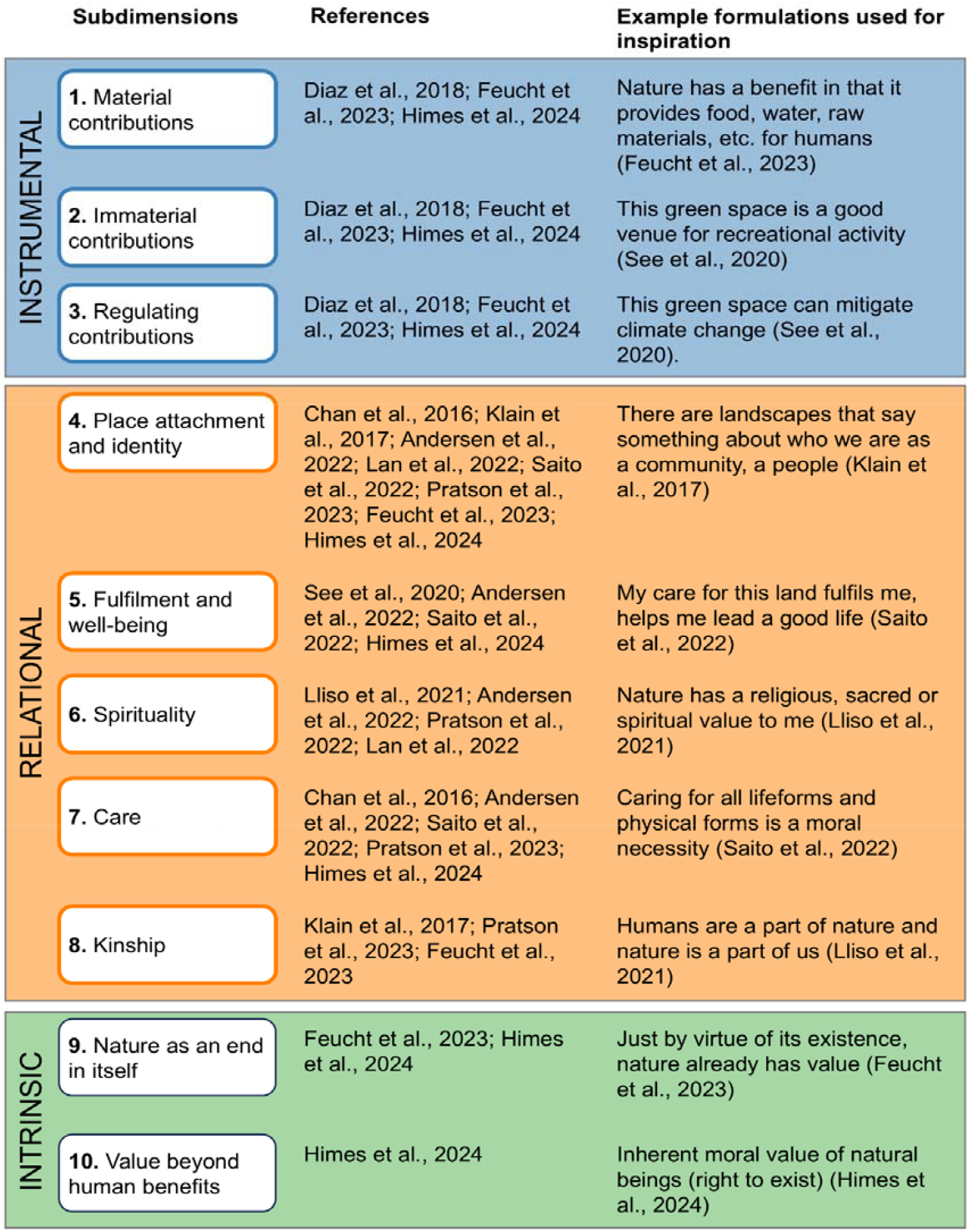
Overview of the values and their subdimensions, including references and examples from previous studies used as inspiration for the PVN.

**Fig. 3:**
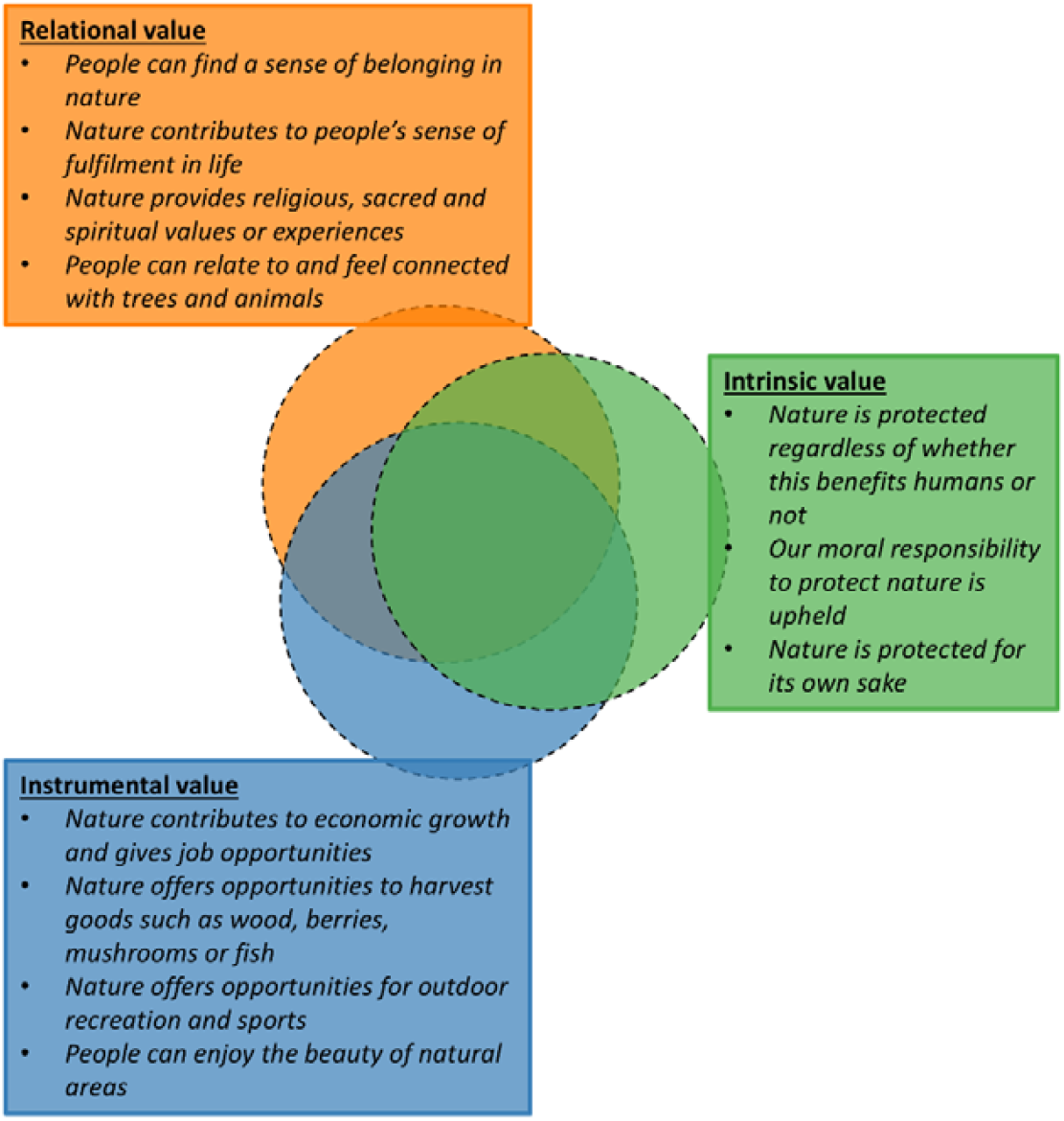
Final items of the Plural Values of Nature (PVN) scale.

As highlighted in recent scholarship (Himes et al., 2024; Kenter et al., 2019) and in the Introduction, the three value dimensions and subdimensions are not mutually exclusive but overlapping with potentially fuzzy boundaries. We therefore expected that some items might display cross-loadings across factors, reflecting the fact that many people endorse multiple value types simultaneously while only a minority endorse values from only one of the three value types. In addition, individual items may load on different (sub)dimensions. Our validation procedure outlined below enabled us to explicitly explore this in the results and reflect on this in the discussion.

The formulation of the items was validated in an online workshop with all coauthors as well as in 16 cognitive interviews with people not professionally engaged in conservation or research in the Netherlands and Denmark. We then pilot tested the 28-items for further refinement and shortening of the scale in representative international samples, resulting in 12 items in the main study and eventually 11 items in the final scale (Fig. 3). Based on the pilot study, the question stem for the final scale was adapted to ‘*Listed below are statements on different aspects that could be important when thinking of nature in the future. Please indicate how much of a priority you feel each aspect should have in* [respondent’s country of residence] *in the future*.’ and responses to each item had to be given on a 7-point scale from ‘not important at all’ to ‘very important’. A full description of all steps and changes in formulations between each step can be found in the supplementary material (S1).

### 2.2 Data Collection

Scale validation was carried out with three independent data collections, each containing samples from multiple countries (Fig. 1). First, an online questionnaire was administered with a convenience sample of university students, some of whom were international students, adding quality to the development of the cross-cultural scale (*n* = 94 & *n* = 101). After further revisions of the scale (see the results section, Table S1, S2, S3), a pilot study was conducted by a professional polling agency, with an online questionnaire targeting a representative sample of the general public in the Netherlands, Spain and Denmark (*n* = 3 x 150) (Table S4). After final iterations of the scale (see results section, Table S1, S5-S8), including a back-translation of the survey to check for translation inconsistencies, it was administered in October 2024 by a professional polling agency as part of a larger online questionnaire on restoration and rewilding. To limit any potential bias related to the aim of the survey, we positioned the items for our PVN scale at the beginning of the questionnaire. The sample consisted of people from the general public in the Netherlands, Spain, Sweden, Poland, and Romania (*n* = 5 x 200) (S9 table). The studies were conducted according to ethical guidelines, including informed consent. The ethics approval for the study was given by the Wageningen Research Ethics Committee for nonmedical studies involving human subjects (WUR-REC-2024-112).

### 2.3 Data Analysis

For the statistical validation of the scale, a range of common scale reliability and validity analysis (Hair et al., 2019) were performed for each of the steps and samples (Fig. 1). Country samples were treated separately in the pretesting and pilot study to support the cross-cultural adaptation and development of the scale. All data analysis was performed in *R* and statistical significance was set at *p* = 0.05.

Across all data collections, we evaluated the performance of individual items using basic descriptive statistics, including mean values, standard deviations (SD), skewness, kurtosis, and response range to assess the clarity of the items and response options and to check for potential ceiling or floor effects (mean values > 6.5 or < 1.5). To assess the appropriateness of our assumed value dimensions, we performed several exploratory and confirmatory factor analysis on our different samples (Fig. 1). In addition, the exploratory analyses provided insights into the degree of overlap between the three value dimensions, and the extent to which they appeared to be distinct from one another in different cultural settings. The reliability of the factors obtained was assessed using Cronbach’s alpha. We also assessed discriminant validity, that is, whether the three value dimensions were sufficiently distinct from one another. Discriminant validity was evaluated using the Fornell–Larcker criterion, which compares the square root of the average variance extracted (AVE) for each factor with the correlations between factors. When inter-factor correlations exceed the square root of AVE, this suggests that dimensions are not fully distinct (Hair et al., 2019). Given our conceptualisation of instrumental, intrinsic, and relational values as overlapping rather than opposing dimensions, we anticipated that issues of discriminant validity might arise. A detailed descriptions of the statistical data analysis in each validation step and for each country, including used configurations, tests for appropriateness, and analysis steps can be found in the Supplementary material S2 Text.

Linear regression models were used to assess the predictive validity of the final scale. Besides the Plural Values of Nature scale, the survey for the final study also contained several items measuring attitudes towards different restoration actions, asking respondents how they would feel about these changes occurring in their country and in an area close to them. We chose four attitude items that asked respondents how they would feel about (1) reducing the influence that humans have on nature at the national (country) level, (2) reducing the influence that humans have on nature at the local (area) level, (3) restoring natural processes and dynamics at the national (country) level, and (4) restoring natural processes and dynamics at the local (area) level. Each of these four items were used as a dependent variable in separate linear regression models, with three composite scales (i.e. instrumental, intrinsic, and relational values) as predictor variables. We then assessed the explained variance of these models.

To illustrate and better understand the plurality of values, we drew inspiration from the Nature-Futures Framework (Kim et al., 2023) and analysed whether people had high endorsement of multiple values simultaneously. We classified respondents according to a simple typology consisting of all possible combinations of the three value dimensions. For example, if people had high endorsement of both instrumental and relational values, they were assigned to the category “Instrumental & Relational” values. This classification was based on respondents’ scores on the composite scales, as these reflect the priority given to different values. Assuming that values above 5 indicate an above-moderate priority (given the 7-point response scale, where 4 corresponds to “moderate priority” and 7 to “highest priority”), we classified respondents into the eight value profiles (Intrinsic; Instrumental; Relational; Intrinsic & Instrumental; Intrinsic & Relational; Instrumental & Relational; Intrinsic & Relational & Instrument; Indifferent).

## 3. Results

After the qualitative pretesting as described in the methods section, we refined and validated the scale in three subsequent quantitative steps in a pre-testing with university students, a pilot survey in three European countries, and a final survey sent to the general public in five European countries. The aim of these steps was to reduce the number of items by focusing the scale on the best performing items and to validate the final scale for its adequacy to measure the three values simultaneously. Here, we report on the results of the final validation test. We will only briefly summarize the results from the pretesting and pilot studies (see S3 Text for extensive description of results) before reporting in-depth about the final validation study.

### 3.1 Quantitative Pre-Testing and Pilot Study

Data analysis of the quantitative pretest (*n* = 94 & *n* =101; see Table S2) revealed a strong ceiling effect (mean scores above 6.5 on a 7-point scale) for some of the items (Table S2). Given that most of the students were enrolled in nature conservation-related programs, we decided not to adjust the items as mean values might be more moderate within more diverse samples. Both samples performed well within the exploratory factor analysis and the identified factor structures generally supported the theoretically expected three-dimensional structure of the scale. In the Danish sample, the three value dimensions emerged as clearly distinct. In contrast, in the Dutch student sample, the item on ‘care’ loaded on the intrinsic values dimension, and the two items related to ‘fulfilment and well-being’ subdimension showed loadings on the instrumental values (Table S3). This points to potential overlaps between relational and intrinsic as well as between relational and instrumental values. Based on the findings of pretesting and additional theoretical reasoning within the research team, a total of eight items were edited in order to improve their clarity and understandability for respondents (Table S1).

The pilot study (*n* = 3 x 150; see Table S4) did not show a strong ceiling effect for any of the items in either sample. Overall, the sample showed statistical adequacy in performing an exploratory factor analysis and the initially retained solution for the complete data set explained 62% of the variation (see Table S6). However, some items, most notably items from the Regulating Benefits subdimension within the instrumental value and the Care subdimension within the relational value caused issues with low variability (e.g. 80% of respondents choosing the two highest response options and/or loaded on a factor associated to one of the other values (S5 Table and S6 Table). The reduction in items resulted in the ‘Regulating Benefits’ and the ‘Care’ subdimension no longer had any related items in the scale, and were therefore removed. We then performed an additional exploratory factor analysis for each country separately (S7 Table), which consistently retained four-factor solutions explaining 64%-69% of the variation. While the general three-dimensional structure was visible, several cross-loadings highlighted the fuzzy boundaries between values and that these might differ across cultural settings. In the Danish sample, the three dimensions again emerged relatively clearly, though one factor contained two of the items from immaterial contributions subdimension together with items from several of the relational values subdimensions. In the Dutch sample, overlaps were more pronounced with one of the factors containing items from all three value dimensions, while the other three factors contained only items from one of the value dimensions. The Spanish sample showed also some cross loadings between items from the instrumental and relational dimension, while intrinsic values build separate factors. Despite some cross loadings, the results supported the overall three-dimensional model, while also illustrating that overlaps between value dimensions incidentally occur, partly dependent of cultural settings. Based on the collective findings and analysis of the quantitative pre-testing and pilot study, we selected 12 of the pilot study items (see items in bold in the Table S7). The grouping of the remaining items according to our theoretical model into instrumental, relational, and intrinsic values dimensions indicated good scale reliability across all countries, suggesting that each items represents its underlying dimension adequately (Cronbach alpha>0.70) (S8 table). See S3 text in the supplementary material for an extensive description of the results. In the hope of increasing the variability in answers, we decided to slightly change the question stem and focus on the priority of values: “*Listed below are statements on different aspects that could be important when thinking of nature in the future. Please indicate how much of a priority you feel each aspect should have in [your country] in the future*” (S1 Table), with a matching answer scale ranging from 1 - Not a priority at all to 4 - Moderate priority and 7 - The highest priority.

### 3.2 Main Survey

The main survey no longer showed strong ceiling effects (S10 Table). However, multivariate normality was not achieved, therefore a robust estimator was used within the confirmatory factor analysis (CFA). Assessment of measurement invariance across countries showed that while configural and metric invariance was established, the scalar invariance was not. This indicates that the factor structure and factor loadings are similar across countries, while there are differences in item intercepts between countries. This might be caused by the cultural differences in response to these items, related to e.g. specific nationally or culturally inspired practices

To make the scale more concise and the number of items balanced across its subdimensions, we decided to reduce the number of items representing immaterial contributions. On the basis of content and theoretical considerations, we decided to remove the item on mental well-being. Doing so results in a similarly good fit of the confirmatory factor analysis (Robust CFI=0.957, Robust TLI=0.942, Robust RMSEA=0.070, SRMR=0.050) and acceptable factor loadings across all countries (Table 1).

**Table 1.**
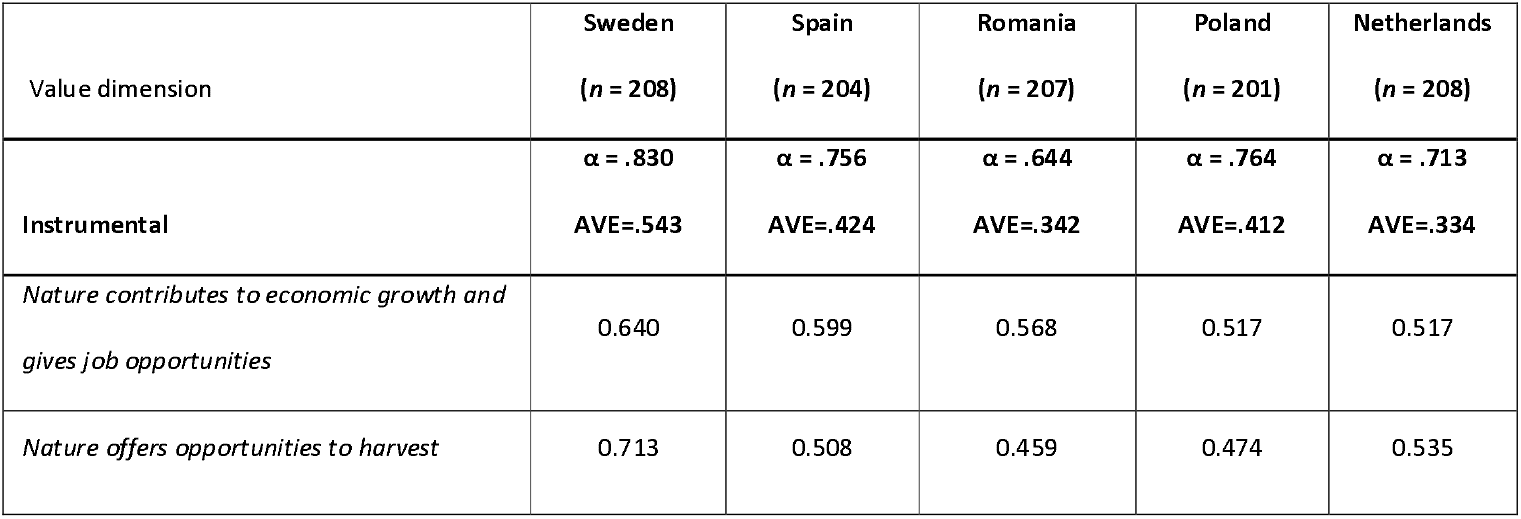
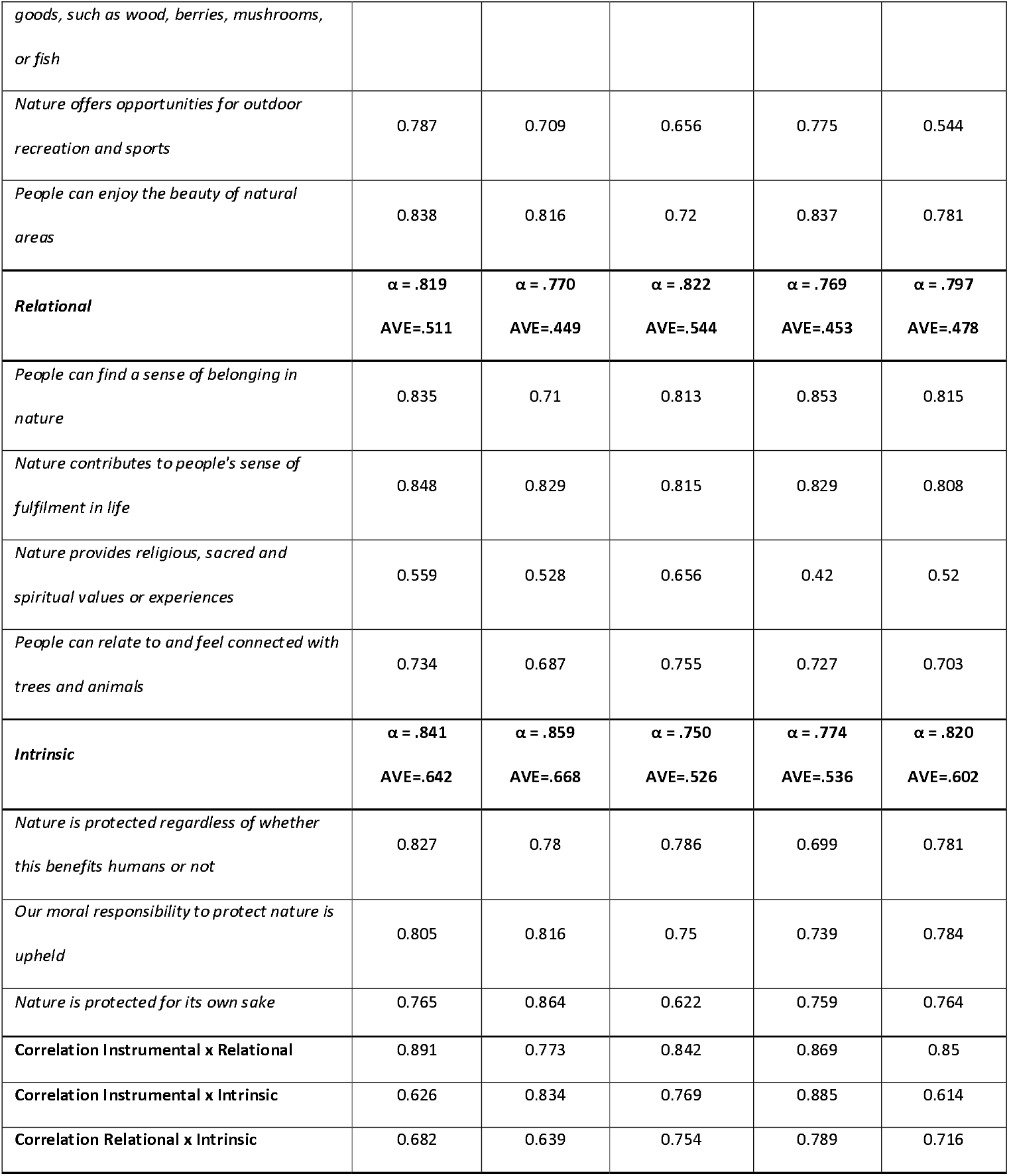
Final Plural Values of Nature scale. Overview of the final items included in the scale, values are standardized factor loadings for the hypothesized factor structure for each of the five samples, including the Cronbach alpha (α) and Average Variance Extracted (AVE) as measures of scale reliability and validity. The last three rows show the correlations between the value dimensions based on the factor analysis.

Concerning scale reliability, the results show acceptable to good internal consistency (Cronbach’s alpha > 0.7) for the three values in all countries, with the exception of *the instrumental value* in Romania (Table 1). In most samples the AVE for *the instrumental and relational value* lies slightly below the recommended threshold of 0.5 indicating weak convergent validity. We assume this is partly due the fact that the items aim to capture different sub-dimensions within the values and thus represent related but distinct facets.

#### Conceptual overlap between values

The confirmatory factor analysis revealed high correlations (r > 0.85) between the three values. Based on follow-up assessment of the Fornell-Larcker criterion, it was confirmed that there seems to be especially a lack of discriminant validity between *instrumental and relational values*. For example, in Sweden, Poland and the Netherlands the correlation between instrumental and relational values exceeded .85, and the AVE for instrumental values fell below .50 (Netherlands AVE = .334; Poland AVE = .412), indicating that these two dimensions are not fully distinct. In Poland, the instrumental and intrinsic dimensions were also highly correlated (r = .885), suggesting fuzziness in distinguishing intrinsic from instrumental values. These patterns are consistent with the expectation that instrumental, relational, and intrinsic values overlap in practice rather than forming sharply distinct categories. This overlap led to the decision to create composite scales before assessing the predictive validity of the different value dimensions.

#### Predictive validity

To assess the predictive validity of our scale and of the three underlying values, we did a linear regression linking the composite scales for each value to four attitudes about nature restoration on the local and national scales that were part of the larger survey. The composite scales showed lower correlations between the three values (r<0.75), allowing us to include them in the same linear regression models while avoiding issues of multicollinearity (VIF<5 for all models). The linear regression models showed a considerable to high predictive validity of the PVN scale (Table 2) with a higher predictive validity toward attitudes at the country level compared to a more local acceptance of restoration actions.

**Table 2.** Predictive validity of the final value scale. Summary of the linear regression models, testing the relationship between value dimensions (independent variables: intrinsic, relational, instrumental) and support for four restoration attitude items (dependent variables) across five countries. Attitude items are stated in the top-left corner of each table panel. The values shown are the standardized regression coefficients (β) and model fit statistics (F statistics with degrees of freedom, adjusted R^2^. Significance Levels: *p < 0.05, **p < 0.01, ***p < 0.001). Bold indicates a significant relationship between the value dimension and attitude item.

| Dependent/<br>independent<br>variable | Sweden | Spain | Romania | Poland | Netherlands |
| --- | --- | --- | --- | --- | --- |
| **Reducing the influence that humans have on nature in the country** | F(3,201)=40.2<br>3,<br>$p < 0.001$ | F(3,199)=17.1<br>2,<br>$p < 0.001$ | F(3,195)=3.3<br>8,<br>$p < 0.05$ | F(3,194)=15.3<br>3,<br>$p < 0.001$ | F(3,198)=19.1<br>8,<br>$p < 0.001$ |
| *Intrinsic* | **0.582\*\*\*** | **0.439\*\*\*** | 0.156 | **0.392\*\*\*** | **0.348\*\*\*** |
| *Relational* | -0.051 | **0.209\*\*** | 0.044 | 0.064 | **0.229\*\*** |
| *Instrumental* | 0.107 | **-0.214\*** | 0.050 | 0.007 | -0.079 |
| **R<sup>2</sup>** | **0.366** | **0.193** | **0.035** | **0.179** | **0.213** |
| **Reducing the influence that humans have on nature in the area** | F(3,201)=7.69,<br>$p < 0.001$ | F(3,199)=5.85,<br>$p < 0.001$ | F(3,195)=2.6<br>3,<br>$p = 0.05$ | F(3,194)=11.4<br>7,<br>$p < 0.001$ | F(3,198)=11.4<br>0,<br>$p < 0.001$ |
| *Intrinsic* | **0.284\*\*\*** | **0.261\*\*** | 0.062 | **0.243\*\*** | **0.235\*\*** |
| *Relational* | 0.136 | **0.190\*** | 0.108 | **0.283\*\*** | **0.263\*\*** |
| *Instrumental* | -0.105 | **-0.236\*** | 0.055 | -0.142 | -0.211** |
| **R<sup>2</sup>** | **0.090** | **0.067** | **0.024** | **0.138** | **0.134** |
| **Restoring natural processes and dynamics in the country** | F(3,201)=33.1<br>2,<br>p<0.001 | F(3,199)=20.4<br>2,<br>p<0.001 | F(3,195)=3.3<br>5,<br>p<0.05 | F(3,194)=27.6<br>7,<br>p<0.001 | F(3,198)=22.7<br>9,<br>p<0.001 |
| *Intrinsic* | **0.557\*\*\*** | **0.414\*\*\*** | 0.076 | **0.429\*\*\*** | **0.383\*\*\*** |
| *Relational* | -0.024 | 0.071 | 0.045 | 0.143 | **0.173\*** |
| *Instrumental* | 0.059 | 0.045 | 0.132 | 0.029 | 0.013 |
| **R2** | **0.321** | **0.224** | **0.034** | **0.289** | **0.245** |
| **Restoring natural processes and dynamics in the area** | F(3,201)=16.7<br>5,<br>p<0.001 | F(3,199)=13.1<br>0,<br>p<0.001 | F(3,195)=6.1<br>7,<br>p<0.001 | F(3,194)=15.3<br>3,<br>p<0.001 | F(3,198)=14.5<br>8,<br>p<0.001 |
| *Intrinsic* | **0.434\*\*\*** | **0.237\*\*** | 0.076 | **0.372\*\*\*** | **0.446\*\*\*** |
| *Relational* | -0.077 | **0.234\*\*** | 0.102 | 0.074 | 0.015 |
| *Instrumental* | 0.100 | -0.005 | 0.186 | 0.025 | -0.094 |
| **R2** | **0.188** | **0.152** | **0.073** | **0.179** | **0.168** |

### 3.3 Plurality of values across study countries

The final scale and the weighted data set indicate in general a high importance (all mean values > 4.5) of the three values in all included countries (Table 3). In Spain and the Netherlands, intrinsic values have the highest mean value, while in Sweden and Romania instrumental values are prioritized the most. Across all countries, the lowest mean value is given to relational values.

**Table 3.** Mean. values for intrinsic, instrumental, and relational values across the five survey countries and the total sample. Composite scale values range from *1 - Not a priority at all* to *4 - Moderate priority* to *7 - The highest priority*. Bold indicates the highest values dimension per country.

|  | Sweden | Spain | Romania | Poland | Netherlands | TOTAL |
| --- | --- | --- | --- | --- | --- | --- |
| **Mean values** |  |  |  |  |  |  |
| *Intrinsic* | 5.72 | **6.25** | 6.06 | **5.96** | **5.70** | **5.93** |
| *Instrumental* | **5.77** | 5.81 | **6.27** | **5.96** | 5.18 | 5.80 |
| *Relational* | 5.07 | 4.86 | 5.86 | 5.32 | 4.77 | 5.17 |

Using our PVN profiles we can illustrate the plurality of values by visualizing the segments of society that prioritize more than one value dimension simultaneously (indicated by an individual expressing a high priority (value >5) on a value type). Figure 4 illustrates the large cooccurrence of the different values across societies, showing that the majority of residents (61.1%-85.4%) in all countries prioritize more than one value type simultaneously. There are, however, differences in the plurality of values (i.e. the prominence of the different value profiles) across the included countries (Fig. 4, S12 Table).

**Fig. 4.**
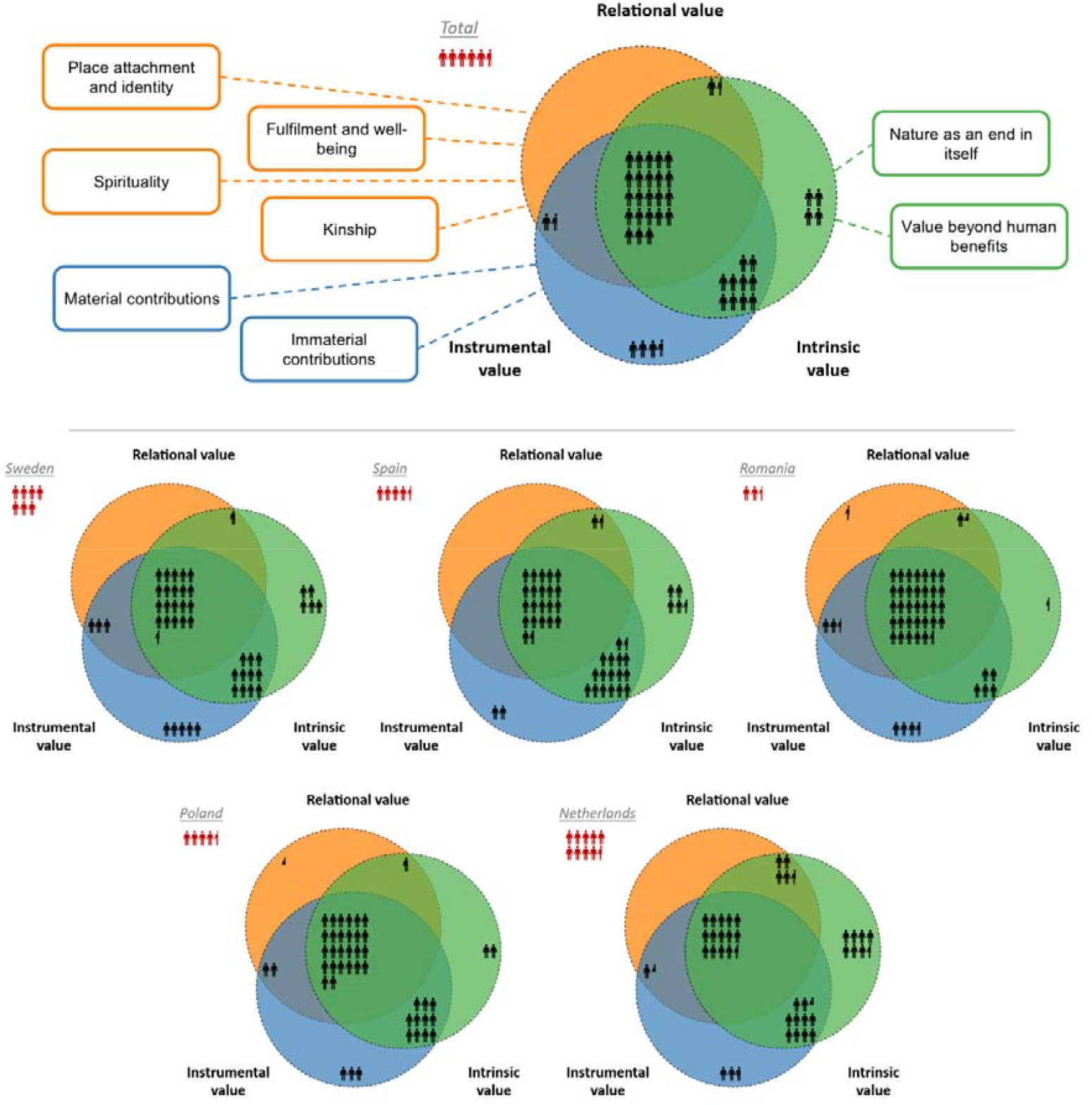
Exploration of plurality of values. Persons indicate the prominence of the different value profiles across the five countries, whereby one person represents 2% of the total population. The red persons outside the circles are those that were classified as none, meaning that they do not express high priority for any of the three values. Figure on top shows average for all five countries (and includes the subdimensions of each value). Figures below represent the five countries separately. (Figure inspired by the Nature Futures Framework; IPBES, 2025).

While individuals expressing high priority for all three value dimensions is the most common profile in all countries the relative share spans from 29.1% in the Netherlands to 67.3% in Romania reflecting a strong pattern of plural value endorsement. The next most common profile in all countries is the combination of intrinsic and instrumental values, ranging from 32.5% in Spain to 10.1% in Romania.

The profiles combining relational with intrinsic and relational with instrumental values are generally less common, with the highest share in The Netherland where 8.9% of respondents simultaneously endorsed relational and intrinsic values. Similarly rare are the ‘pure’ profiles meaning segments of respondent that only prioritize one of the value dimensions. While 14.8% of Dutch respondents prioritized only intrinsic values and 10.6% of Swedish respondents prioritized purely instrumental values, respondents prioritizing relational values alone were missing in three out of the five countries. Even in Romania & Poland, they comprised only 0.5-1.0% of the population. Taken together, these patterns highlight that ‘clear-cut’ positions (endorsing only one value type) are relatively rare, while plural combinations dominate across all five countries.

To examine the relevance of the plurality of values in predicting support for conservation measures, we plotted the value profiles (collectively for all samples) against the four attitude items (Fig 5). This did not only reaffirm the relevance (i.e. predictive validity) of the scale but also showed considerable differences among the value profiles. Those expressing no priority for any value (i.e. ‘Indifferent’) consistently show the lowest support for restoration actions. Interestingly, the relative position of the ‘Indifferent’ group and the distance to the other value profiles shifts depending on scale. Asking about the national level, they differ considerably from the other profiles. Regarding restoration actions at the local level (in the area where people live), they align more closely with the purely instrumental group and the ‘instrumental & relational’ respondents. In contrast, people prioritizing intrinsic & relational values or all three values simultaneously show the highest levels of support. Importantly, respondents with plural combinations of values (e.g., instrumental & relational, or all three together) generally expressed stronger support than those with single value orientations, underscoring the role of plurality in motivating restoration support.

**Fig 5.**
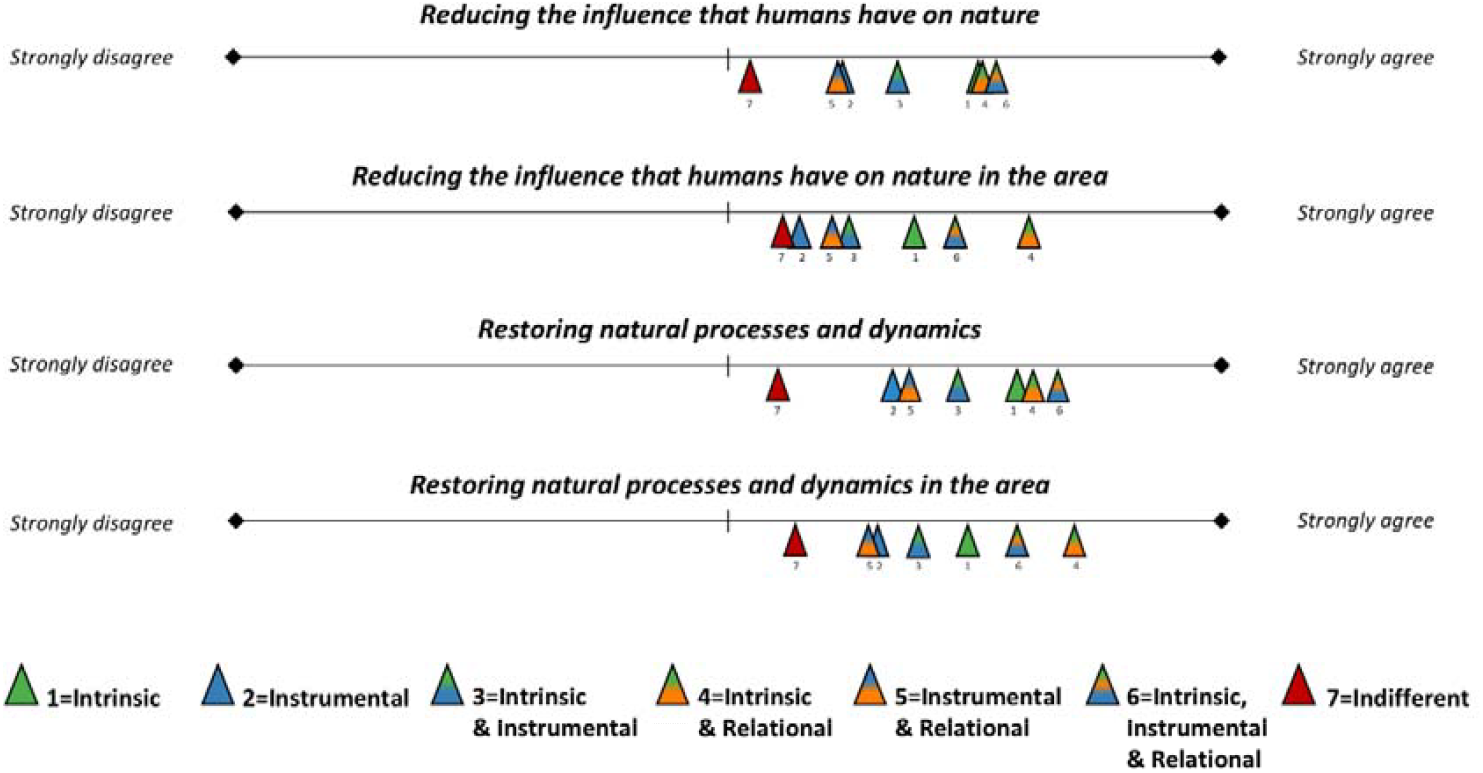
Mean support for restoration actions (four attitude items) across value typologies. The lines represent a 7-point Likert scale (left end point = strongly disagree, right endpoint = strongly agree), on which the typologies are placed according to their mean score across the total sample. (Note that due to the small number of participants falling purely within the relational value, this type was not plotted for the attitude scales).

## 4. Discussion

The presented Plural Value of Nature (PVN) scale is a new psychometric scale and survey-based social indicator to measure people’s value of nature, differentiating between relational, intrinsic, and instrumental values. The PVN is the first cross-nationally validated scale that captures relational, intrinsic, and instrumental values simultaneously. The scale recognises several things: the plurality of values held across societies; that individuals can endorse values of nature related to the intrinsic, instrumental and relational value types simultaneously; and that values may be prioritized differently between people and within and between countries, leading to potential differences in nature-related preferences, attitudes, and behaviours. Cross-national validation studies in five countries (*n* = 1028) show internal consistency and reliability of the scale. In addition, the scale has considerable to high predictive validity, explaining up to 37 % of variability in attitudes towards ecological restoration. In particular, intrinsic values were found to have especially strong predictive value. These results confirm the validity and practical value of the scale, and its usefulness across different national contexts within Europe. Initial results show that the majority of people in all countries tested, assigned a moderate to high priority (values above 4 on a 7-point scale) to *all* three value types (Table 3), confirming recent calls to recognize not only instrumental and intrinsic values but also relational nature values (Andersen et al., 2022; Chan et al., 2016). The results can be seen as strong empirical confirmation of the need to focus on the plurality of values. Interestingly, our analysis also shows that this plurality of values also impacts people’s attitude towards nature restoration. People that endorse multiple value show more positive attitudes towards nature restoration than people endorsing only one specific value (either instrumental, intrinsic *or* relational value), with people who endorse all three values showing the highest support for such actions. While relational values also had positive correlations to restoration attitudes, support for the relational value was on average lower in all test countries, most notably for the item on the religious, sacred, and spiritual values of nature. This may be a reflection of our focus on the general public of European countries, where spiritual values may be less prominent than in e.g., indigenous communities (Verschuuren & Brown, 2019). Interestingly, the impact of values shifts depending on spatial scale (Table 2), suggesting that instrumental considerations may dominate in national debates, whereas relational ties to place and community become more influential when people think about ecological restoration in local contexts.

### 4.1 Plurality of values and scale development challenges

In developing a scale to measure the values of nature, it is important to reflect on epistemological and ontological discussions regarding the nature of the three values (Kenter et al., 2019), most notably of relational values (Gould et al., 2024; Stålhammar & Thorén, 2019), and on the potential fuzzy boundaries between these values (Himes et al., 2024; Kenter et al., 2019). Instrumental and intrinsic values are well established in literature, whereas the conceptual debate on relational values, and its relationship to the two other values, is ongoing (Ibid.). This conceptual challenge was reflected in the process of conceptualising and operationalising relational values for our scale. Validation of the scale showed that fuzzy boundaries do indeed exist between the values. Most notably, our results show that especially the instrumental and relational dimensions are closely related and not fully distinct. Difficulty in delineating between relational and instrumental values was a persistent issue, from the initial pre-testing to the final scale. This, however, was not unexpected, as in theory relational values are described as “anthropocentric yet non-instrumental”, and thus closely related to, or overlap with, at least a subsection of instrumental values (Himes & Muraca, 2018). We encountered overlaps such as this when designing the scale. For example, only after a thorough review of the classifications of beauty or spirituality in the literature into different value types (or into two or three types simultaneously) (e.g. Anderson et al., 2022; R. Gould et al., 2024; Himes et al., 2024; James, 2019), and after internal discussions and cognitive interviews, we decided to conceptualize the spiritual component of nature as a relational value and the beauty of nature as an instrumental value and tried to formulate the items in line with this (partial) conceptualisation (see also Table S1 for these changes throughout the process). While confirmatory factor analysis confirmed most respondents indeed interpreted these items as respectively relational and instrumental values, analysis also showed that at the national level the items sometimes also scored on other value types. In addition, in several countries, the item on berry picking - conceptualised as an instrumental value by the researchers - also loaded on to the factor related to relational values. This again is an example of conceptual overlap, where in some targeted countries (e.g., Denmark) berry picking is also a culturally significant practice for relating to nature and thus is not only valued for the instrumental value of the berries themselves. Furthermore, instrumental items reflecting Nonmaterial Contributions to People (NCP’s) - or cultural ecosystem services in ESS terminology - showed loadings on both the relational and instrumental value factor. Consequently, approaches to operationalise instrumental values through the concept of ecosystem services may need to be reconsidered in light of the more recent debate on relational values (e.g. Plieninger et al., 2013). In general, as highlighted by literature on relational values, Nature’s Contributions to People, or ecosystem services should not be considered as one-directional contributions, but as multi-directional relationships between humans and nonhuman nature, where the relationship itself holds value and contributes to nature and people alike (Himes et al., 2024; West et al., 2018).

More generally, the overlap and fuzzy boundaries between the values raise the question of whether developing such a psychometric scale is feasible at all, or whether it would be better to use three independent scales for capturing intrinsic, instrumental, and relational values. While this may seem more conceptually valid, we argue that while such separate scales would suggest conceptual clarity for each value type – capturing better the full complexity of the values – they would be unable to explicitly explore the overlaps and boundaries between the values. In our view, combining all three values is crucial to understanding and exploring the plurality of values and the fuzzy boundaries between them. Obviously, developing a combined scale means compressing meaning, reducing complexity – and, not least, risking validity. Although the reduction of complexity in this scale may not fit the potential use for in-depth analysis of each individual value, its feasibility should be seen in relation to its practical use – as one of the criteria we set up for the scale development – and so, as Stern et al. (1998) argue in their seminal paper, it is crucial to develop a brief ‘inventory’ of values that is suitable for use in survey research, as longer instruments are simply impractical – not least in relation to challenges of survey fatigue (Jeong et al., 2023). Based on experience with e.g. the 15-item NEP-scale (Dunlap et al., 2000), we argue an 11-item scale is a feasible length to add to question batteries in environmental research.

### 4.3 Potential limitations

Three potential limitations of PVN should be discussed. First, quantitatively measuring values of nature is a challenging task. The measurement of relational values is especially challenging, not only because of epistemological and conceptual differences between different schools of thought, but also because they relate to a wide variety of relationships – both relationships between people and nature and nature-meditated relationships between people and people (Chan et al., 2016). While we explored our experiences and empirical findings in relation to some of these conceptual challenges above, we do not engage in-depth with these issues which have been explored elsewhere (e.g. Rawluk et al., 2019). Meanwhile, we agree with Schultz and Martin-Ortega (2018), who argue that exploring both quantitative and qualitative approaches to the understanding of relational values is helpful for our understanding of values – and, we would add, of the plurality of values. In addition, Jacobs et al (2018) argue that socio-cultural valuation methods, such as surveys, are better suited than other elicitation methods to capture such values and should thus be developed further. Further developing quantitative approaches to understand the plurality of values of nature not only expands the evidence base for the relevance of acknowledging diverse values in society but can also help contribute to a better understanding of potential overlaps and divergence between values. In addition, it may contribute to more interdisciplinary research, linking more quantitative fields, such as environmental psychology, to more qualitative fields, such as ethnography or critical studies. Finally, it may contribute to the political legitimacy of environmental decision-making, where numbers and statistics may have more discursive power than more qualitative and interpretive approaches (Schulz & Martin-Ortega, 2018).

Secondly, one could ask whether it is possible to measure values without explicit reference to specific places. We believe that with the formulation set out in the questionnaire (“…considering nature *in your country*…”), it does stimulate respondents to link the values to a (more specific) spatial context – that is, their national context. That people care about nature at a national level is shown in multiple contexts (Bakhtiari et al., 2018), and hence we would also expect values to be relevant here. In addition, the cognitive interviews during semantic validation suggested people were quite able and willing to reflect on items related to, e.g., recreation, spirituality and kinship *without* reference to specific, well-defined places. Here, our proposed scale may be situated in-between existing traditions, ranging from the usually highly contextualized sense-of-place literature (Williams & Vaske, 2003) (but see Raymond et al., 2023 for more fluid approaches to senses of place), to generic approaches trying to gauge overall connection to, concern for, or affinity to nature (e.g. Dunlap et al., 2000; Mayer & Frantz, 2004). Meanwhile, researchers who wish to focus on values related to local or regional nature projects can relatively easy do so by adapting the stem of the scale (e.g., ‘…considering nature in your region’).

Thirdly to maximise the usefulness of the scale for transformative practices and transformative science, we choose to focus on ‘future nature’ when asking about values of nature. Here, one can ask what the respondents will perceive - how far into the future will that be? This did not arise as a matter of ambiguity or concern during the scale validation process. Again, as time spans differ in terms of changes in nature (e.g., a forest can take more than 100 years to fully grow up, whilst changes in meadows or moors can be considerable after maybe ten or twenty years), applying a restricted time length could again act as too much of a restriction. Obviously, it still opens the question of what period respondents had in mind when answering the survey, and if a specific time length is in question, it may just be useful to specify it and to adapt the scale to be used in such a manner.

### 4.4 The PVN as a survey-based indicator of plural values

The focus of our study was on developing and validating the PVN, not on measuring the relative support for each of the values in the five countries. However, results do suggest that not only instrumental and intrinsic but also relational values are recognized by the general public. Recognising multiple values is argued in the literature to offer powerful pathways to reconnect people to their natural environment (Beery et al., 2023; Richardson et al., 2020). In addition, elucidating nature-positive values is seen as an important leverage point for transformations (Horcea-Milcu et al., 2023). Indeed, as argued above, our results suggest that mobilising multiple values simultaneously may strengthen support for conservation practices.

We consider PVN a a survey-based social indicator that can help to take stock of the current values that societies have regarding the future development of nature, and as a monitoring tool to observe potential shifts in specific societal values of nature over time. In this sense, the PVN can be understood as a survey-based social indicator of plural values of nature. Rather than measuring ecological status directly, it captures a societal condition that is increasingly recognised as relevant for biodiversity governance and sustainability transformations: the extent to which different publics prioritise instrumental, intrinsic, and relational values of nature (Pascual et al., 2023). As such, the PVN complements biophysical indicators by providing a concise and repeatable measure of the values that may shape public support for conservation, restoration, and nature-positive transformation pathways.

We discussed that the PVN has strong explanatory power for people’s attitudes towards ecological restoration. Based on this predictive validity, the PVN could also be used to explore people’s preferences for future developments impacting nature, including approaches for transformative change. Using the PVN in transformative dialogues and processes could contribute to the inclusiveness of these processes. Here, the link to the Nature Futures Framework can be relevant, an approach to develop scenarios for, what is stated as, nature-positive futures (Pereira et al., 2020), recognising the diversity and - more recently - the plurality of values of nature (Durán et al., 2023). In situations where qualitative approaches may not be possible (such as the number of stakeholders being too high, or trust and engagement with stakeholders may be too low), using the PVN in surveys can contribute to more inclusive and just transformation process.

## 5. Conclusions

Our aspiration for this research was to develop a a concise, validated measure of the co-occurrence of multiple values of nature: the Plural Values of Nature (PVN) scale. By providing a repeatable survey-based measure, the PVN can function as a social indicator for monitoring plural values of nature across populations and over time. This indicator function is especially relevant in contexts where biodiversity and restoration policies require not only ecological monitoring, but also an understanding of the societal values that shape support, contestation and implementation.

Such quantification can be used to increase our understanding of the plural values of nature and link it to elicited attitudes towards, concrete management initiatives across stakeholder groups and different population segments. Moreover, such a scale may help to increase understanding of the plurality of values and can be used to assess the endorsement of different values of nature across regions, cultures, countries, and time-periods. Based on the strong predictive power, we expect that the scale can contribute to our understanding of attitudes towards preservation, conservation and restoration of nature. We also hope that its use can facilitate more inclusive dialogues for effective and just transformative change, by sensitising change efforts to the plural values of diverse communities and informing policies on smaller or larger geographical scales.

The scale developed is simple and short – so that it can be of practical use in surveys linked to other research questions being raised, where survey length and fatigue are common concerns. We have shown that the PVN works in different countries within Europe. Testing the usefulness of the scale in different cultures and beyond Europe will be needed to expand its value to other countries in the Global North. We have found that the scale, despite its chosen simplicity, is able to identify intrinsic, relational and instrumental values for nature held by the general public. We find some overlap between the value categories, likely reflecting inherent elements of fuzziness in the definition as well as in the empirical specification of these values. We acknowledge that brevity of the scale comes at a cost – there may be nuances of the plurality that we miss, despite a thorough empirical approach. We encourage future research to elaborate further on these potential limitations and explore use of the scale beyond the European context and, indeed, explore the use of the scale beyond a European context.

## Ethical approval

Research has been done according to ethical standards for human subject research, including formal approval of the study by the appropriate institutional ethics commission. Ethics committee: WUR Research Ethics Commission for review of nonmedical studies with human subjects (WUR-REC) APPROVAL NUMBER: 2024-112 FORM OF CONSENT: written consent.

## Acknowledgements

This manuscript and the underlying research received financial support from the European Union’s Horizon Europe research and innovation program under grant agreement 101081251 (wildE). Views and opinions expressed are, however, those of the authors only and do not necessarily reflect those of the European Union. Neither the European Union nor the granting authority can be held responsible for them.

## Data availability statement

We intend to archive our data in the Data Station for Social Sciences and Humanities (DANS) upon acceptance.

## Declaration of generative AI and AI-assisted technologies in the manuscript preparation process

During the preparation of this work, the author(s) used ChatGPT for language refinement of author-provided text, including grammar, sentence structure, clarity, and readability. The tool was not used to generate original scientific content, analyse data, or draw conclusions. After using this tool, the author(s) reviewed and edited the content as needed and take full responsibility for the content of the published article.

## References

Andersen, G., Fløttum, K., Carbou, G., & Gjesdal, A. M. (2022). People’s Conceptions and Valuations of Nature in the Context of Climate Change [Article]. Environmental Values, 31(4), 397–420. 10.3197/096327121X16328186623850

Anderson, C. B., Athayde, S., Raymond, C. M., Vatn, A., Arias, P., & Gould, R. K., Kenter, J., Muraca, B., Sachdeva, S., Samakov, A., Zent, E., Lenzi, D., Murali, R., Balvanera, P., Pascual, U., Christie, M., Baptiste, B., and González-Jiménez, D.. (2022). Chapter 2: Conceptualizing the diverse values of nature and their contributions to people. In P. Balvanera, Pascual, U., Christie, M., Baptiste, B., and González-Jiménez, D. (Ed.), Methodological Assessment Report on the Diverse Values and Valuation of Nature of the Intergovernmental Science-Policy Platform on Biodiversity and Ecosystem Services. IPBES secretariat. 10.5281/zenodo.6493134

Bakhtiari, F., Jacobsen, J. B., Thorsen, B. J., Lundhede, T. H., Strange, N., & Boman, M. (2018). Disentangling Distance and Country Effects on the Value of Conservation across National Borders. Ecological Economics, 147, 11–20. 10.1016/j.ecolecon.2017.12.019

Beery, T., Stahl Olafsson, A., Gentin, S., Maurer, M., Stålhammar, S., Albert, C., Bieling, C., Buijs, A., Fagerholm, N., Garcia-Martin, M., Plieninger, T., & M. Raymond, C. (2023). Disconnection from nature: Expanding our understanding of human–nature relations. People and Nature, 5(2), 470–488. 10.1002/pan3.10451

Bennett, N. J., Blythe, J., Cisneros-Montemayor, A. M., Singh, G. G., & Sumaila, U. R. (2019). Just transformations to sustainability [Review]. Sustainability (Switzerland), 11(14), Article 3881. 10.3390/su11143881

Boateng, G., Neilands, T., Frongillo, E., Melgar-Quinonez, H., & Young, S. (2018). Best Practices for Developing and Validating Scales for Health, Social, and Behavioral Research: A Primer. Front Public Health, 6, 149. 10.3389/fpubh.2018.00149

Boateng, G. O., Neilands, T. B., Frongillo, E. A., Melgar-Quiñonez, H. R., & Young, S. L. (2018). Best Practices for Developing and Validating Scales for Health, Social, and Behavioral Research: A Primer [Review]. Frontiers in Public Health, 6. 10.3389/fpubh.2018.00149

Brown, T. C. (1984). The concept of value in resource allocation. Land Economics, 60(3), 231–246. 10.2307/3146184

Buijs, A., Hoogstra-Klein, M., de Boer, T., Dressel, S., & Langers, F. (2025). Evidence for increasing public support for nature conservation: A 15-year longitudinal analysis of nature conservation attitudes in the Netherlands. Biological Conservation, 308. 10.1016/j.biocon.2025.111239

Buijs, A. E. (2009). Lay people’s images of nature: frameworks of values, beliefs and value orientations. Society and Natural Resources, 22(5), 417–432.

Buijs, A. E., Elands, B. H. M., & Langers, F. (2009). No wilderness for immigrants: Cultural differences in images of nature and landscape preferences. Landscape and Urban Planning, 91(3), 113–123. 10.1016/j.landurbplan.2008.12.003

Chan, K. M. A., Balvanera, P., Benessaiah, K., Chapman, M., Díaz, S., Gómez-Baggethun, E., Gould, R., Hannahs, N., Jax, K., Klain, S., Luck, G. W., Martín-López, B., Muraca, B., Norton, B., Ott, K., Pascual, U., Satterfield, T., Tadaki, M., Taggart, J., & Turner, N. (2016). Why protect nature? Rethinking values and the environment. Proceedings of the National Academy of Sciences of the United States of America, 113(6), 1462–1465. 10.1073/pnas.1525002113

Chan, K. M. A., Gould, R. K., & Pascual, U. (2018). Editorial overview: Relational values: what are they, and what’s the fuss about? Current Opinion in Environmental Sustainability, 35, A1–A7. 10.1016/j.cosust.2018.11.003

Diaz, S., Pascual, U., Stenseke, M., Martin-Lopez, B., Watson, R. T., Molnar, Z., Hill, R., Chan, K. M. A., Baste, I. A., Brauman, K. A., Polasky, S., Church, A., Lonsdale, M., Larigauderie, A., Leadley, P. W., van Oudenhoven, A. P. E., van der Plaat, F., Schroter, M., Lavorel, S., Shirayama, Y. (2018). Assessing nature’s contributions to people. Science, 359(6373), 270–272. 10.1126/science.aap8826

Dunlap, R. E., Van Liere, K. D., Mertig, A. G., & Jones, R. E. (2000). Measuring endorsement of the new ecological paradigm: A revised NEP scale. The Journal of social issues, 56(3), 425–442. http://sfx.library.wur.nl:9003/sfx_local?sid=SP%3APSYI;id=pmid%3A;id=doi%3A10.1111%2F0022-4537.00176;issn=0022-4537;isbn=;volume=56;issue=3;spage=425;pages=425-442;date=2000;title=Journal%20of%20Social%20Issues;atitle=Measuring%20endorsement%20of%20the%20new%20ecological%20paradigm%3A%20A%20revised%20NEP%20scale.;aulast=Dunlap;pid=%3CAN%3E2001-14019-004%3C%2FAN%3E%3CAU%3EDunlap%2C%20Riley%20E%3BVan%20Liere%2C%20Kent%20D%3BMertig%2C%20Angela%20G%3BEmmet%20Jones%2C%20Robert%3C%2FAU%3E%3CDT%3EJournal%3BPeer%20Reviewed%20Journal%3BEmpirical%20Study%3C%2FDT%3E

Durán, A. P., Kuiper, J. J., Aguiar, A. P. D., Cheung, W. W. L., Diaw, M. C., Halouani, G., Hashimoto, S., Gasalla, M. A., Peterson, G. D., Schoolenberg, M. A., Abbasov, R., Acosta, L. A., Armenteras, D., Davila, F., Denboba, M. A., Harrison, P. A., Harhash, K. A., Karlsson-Vinkhuyzen, S., Kim, H. J., Pereira, L. M. (2023). Bringing the Nature Futures Framework to life: creating a set of illustrative narratives of nature futures [Article]. Sustainability Science. 10.1007/s11625-023-01316-1

Feucht, V., Dierkes, P. W., & Kleespies, M. W. (2023). The different values of nature: a comparison between university students’ perceptions of nature’s instrumental, intrinsic and relational values. Sustainability Science, 18(5), 2391–2403. 10.1007/s11625-023-01371-8

Gould, R., Muraca, B., Himes, A., & Hackenburg, D. (2024). Biodiversity and Relational Values. In Encyclopedia of Biodiversity (pp. 8–17). 10.1016/b978-0-12-822562-2.00091-8

Gould, R. K., Himes, A., Anderson, L. M., Arias Arévalo, P., Chapman, M., Lenzi, D., Muraca, B., & Tadaki, M. (2024). Building on Spash’s critiques of monetary valuation to suggest ways forward for relational values research. Environmental Values, 33(2), 139–162.

Hair, J. F., Gabriel, M. L. D. S., da Silva, D., & Braga Junior, S. (2019). Development and validation of attitudes measurement scales: fundamental and practical aspects. RAUSP Management Journal, 54(4), 490–507. 10.1108/RAUSP-05-2019-0098

Himes, A., & Muraca, B. (2018). Relational values: the key to pluralistic valuation of ecosystem services. Current Opinion in Environmental Sustainability, 35, 1–7. 10.1016/j.cosust.2018.09.005

Himes, A., Muraca, B., Anderson, C. B., Athayde, S., Beery, T., Cantu-Fernandez, M., Gonzalez-Jimenez, D., Gould, R. K., Hejnowicz, A. P., Kenter, J., Lenzi, D., Murali, R., Pascual, U., Raymond, C., Ring, A., Russo, K., Samakov, A., Stalhammar, S., Thoren, H., & Zent, E. (2024). Why nature matters: A systematic review of intrinsic, instrumental, and relational values. BioScience, 74(1), 25–43. 10.1093/biosci/biad109

Horcea-Milcu, A. I., Koessler, A. K., Martin, A., Rode, J., & Moreno Soares, T. (2023). Modes of mobilizing values for sustainability transformation [Review]. Current Opinion in Environmental Sustainability, 64, Article 101357. 10.1016/j.cosust.2023.101357

Hughes, A. C., & Grumbine, R. E. (2023). The Kunming-Montreal Global Biodiversity Framework: what it does and does not do, and how to improve it. Frontiers in Environmental Science, 11. 10.3389/fenvs.2023.1281536

IPBES. (2022). Summary for Policymakers of the methodological Assessment Report on the Diverse Values and Valuation of Nature of the Intergovernmental Science-Policy Platform on Biodiversity and Ecosystem Services.

IPBES. (2024). Thematic Assessment Report on the Underlying Causes of Biodiversity Loss and the Determinants of Transformative Change and Options for Achieving the 2050 Vision for Biodiversity.

IPBES (2025). The Nature Futures Framework, a flexible tool to support the development of scenarios and models 592 of desirable futures for people, nature and Mother Earth, and its methodological guidance, version Dec 593 2025. IPBES secretariat, Bonn, Germany.

Jacobs, S., Martín-López, B., Barton, D. N., Dunford, R., Harrison, P. A., Kelemen, E., Saarikoski, H., Termansen, M., García-Llorente, M., Gómez-Baggethun, E., Kopperoinen, L., Luque, S., Palomo, I., Priess, J. A., Rusch, G. M., Tenerelli, P., Turkelboom, F., Demeyer, R., Hauck, J., Smith, R. (2018). The means determine the end – Pursuing integrated valuation in practice. Ecosystem Services, 29, 515–528. 10.1016/j.ecoser.2017.07.011

James, S. P. (2019). Natural meanings and cultural values [Article]. Environmental Ethics, 41(1), 3–16. 10.5840/enviroethics20194112

James, S. P. (2022). Against Relational Value. The Harvard review of Philosophy, 29, 45–54.

Jeong, D., Aggarwal, S., Robinson, J., Kumar, N., Spearot, A., & Park, D. S. (2023). Exhaustive or exhausting? Evidence on respondent fatigue in long surveys [Article]. Journal of Development Economics, 161, Article 102992. 10.1016/j.jdeveco.2022.102992

Kempton, W., Boster, J. S., & Hartley, J. A. (1995). Environmental Values in American Culture. MIT Press.

Kenter, J. O., Raymond, C. M., van Riper, C. J., Azzopardi, E., Brear, M. R., Calcagni, F., Christie, I., Christie, M., Fordham, A., Gould, R. K., Ives, C. D., Hejnowicz, A. P., Gunton, R., Andra-Ioana, H.-M., Kendal, D., Kronenberg, J., Massenberg, J. R., Seb, O. C., Ravenscroft, N., Thankappan, S. (2019). Loving the mess: navigating diversity and conflict in social values for sustainability. Sustainability Science, 14(5), 1439–1461. 10.1007/s11625-019-00726-4

Kim, H., Peterson, G. D., Cheung, W. W. L., Ferrier, S., Alkemade, R., Arneth, A., Kuiper, J. J., Okayasu, S., Pereira, L., Acosta, L. A., Chaplin-Kramer, R., den Belder, E., Eddy, T. D., Johnson, J. A., Karlsson-Vinkhuyzen, S., Kok, M. T. J., Leadley, P., Leclère, D., Lundquist, C. J., Pereira, H. M. (2023). Towards a better future for biodiversity and people: Modelling Nature Futures. Global Environmental Change, 82. 10.1016/j.gloenvcha.2023.102681

Klain, S. C., Olmsted, P., Chan, K. M. A., & Satterfield, T. (2017). Relational values resonate broadly and differently than intrinsic or instrumental values, or the New Ecological Paradigm. PLoS ONE, 12(8), e0183962. 10.1371/journal.pone.0183962

Kloek, M. E., Buijs, A. E., Boersema, J. J., & Schouten, M. G. C. (2017). Beyond Ethnic Stereotypes – Identities and Outdoor Recreation Among Immigrants and Nonimmigrants in the Netherlands. Leisure Sciences, 39(1), 59–78. 10.1080/01490400.2016.1151843

Lengieza, M. L., Aviste, R., & Swim, J. K. (2023). Nature as community: An overlooked predictor of proenvironmental intentions. Journal of Environmental Psychology, 91. 10.1016/j.jenvp.2023.102127

Li, J., Warchold, A., & Pradhan, P. (2025). Revisiting social foundations and well-being indicators for sustainability: Insights from a systematic literature review. Ecological Indicators, 178. 10.1016/j.ecolind.2025.113890

Lockwood, M. (1999). Humans Valuing Nature: Synthesising Insights from Philosophy, Psychology and Economics. Environmental Values, 8(3), 381. http://www.swetswise.com/link/access?db?issn=0963-2719&vol=00008&iss=00003&year=1999&page=381&ft=1

Lou, X., Mei, D., & Li, L. M. W. (2025). Development and validation of a scale measuring intrinsic, relational, and instrumental environmental values. Journal of Environmental Psychology, 106. 10.1016/j.jenvp.2025.102744

Luque-Lora, R. (2023). The Trouble with Relational Values. Environmental Values, 32(4), 411–431. 10.3197/096327122x16611552268681

Mayer, F. S., & Frantz, C. M. (2004). The connectedness to nature scale: A measure of individuals’ feeling in community with nature. Journal of Environmental Psychology, 24(4), 503–515. 10.1016/j.jenvp.2004.10.001

Mrotek, A., Anderson, C. B., Valenzuela, A. E. J., Manak, L., Weber, A., Van Aert, P., Malizia, M., & Nielsen, E. A. (2019). An evaluation of local, national and international perceptions of benefits and threats to nature in Tierra del Fuego National Park (Patagonia, Argentina). Environmental Conservation, 46(4), 326–333. 10.1017/s0376892919000250

Muraca, B. (2011). The map of moral significance: A new axiological matrix for environmental ethics. Environmental Values, 20(3), 375–396.

Norton, B., & Sanbeg, D. (2021). Relational values: a unifying idea in environmental ethics and evaluation? Environmental Values, 30(6), 695–714.

Pascual, U., Balvanera, P., Anderson, C. B., Chaplin-Kramer, R., Christie, M., González-Jiménez, D., Martin, A., Raymond, C. M., Termansen, M., Vatn, A., Athayde, S., Baptiste, B., Barton, D. N., Jacobs, S., Kelemen, E., Kumar, R., Lazos, E., Mwampamba, T. H., Nakangu, B., Zent, E. (2023). Diverse values of nature for sustainability. Nature. 10.1038/s41586-023-06406-9

Pascual, U., Balvanera, P., Díaz, S., Pataki, G., Roth, E., Stenseke, M., Watson, R. T., Başak Dessane, E., Islar, M., Kelemen, E., Maris, V., Quaas, M., Subramanian, S. M., Wittmer, H., Adlan, A., Ahn, S., Al-Hafedh, Y. S., Amankwah, E., Asah, S. T., Yagi, N. (2017). Valuing nature’s contributions to people: the IPBES approach. Current Opinion in Environmental Sustainability, 26-27, 7–16. 10.1016/j.cosust.2016.12.006

Patterson, J., Schulz, K., Vervoort, J., van der Hel, S., Widerberg, O., Adler, C., Hurlbert, M., Anderton, K., Sethi, M., & Barau, A. (2017). Exploring the governance and politics of transformations towards sustainability. Environmental Innovation and Societal Transitions, 24, 1–16. 10.1016/J.EIST.2016.09.001

Pereira, L. M., Davies, K. K., den Belder, E., Ferrier, S., Karlsson-Vinkhuyzen, S., Kim, H., Kuiper, J. J., Okayasu, S., Palomo, M. G., Pereira, H. M., Peterson, G., Sathyapalan, J., Schoolenberg, M., Alkemade, R., Carvalho Ribeiro, S., Greenaway, A., Hauck, J., King, N., Lazarova, T., Lundquist, C. J. (2020). Developing multiscale and integrative nature–people scenarios using the Nature Futures Framework. People and Nature, 2(4), 1172–1195. 10.1002/pan3.10146

Plieninger, T., Dijks, S., Oteros-Rozas, E., & Bieling, C. (2013). Assessing, mapping, and quantifying cultural ecosystem services at community level. Land Use Policy, 33, 118–129. 10.1016/j.landusepol.2012.12.013

Pratson, D. F., Adams, N., & Gould, R. K. (2023). Relational values of nature in empirical research: A systematic review. People and Nature, 5(5), 1464–1479. 10.1002/pan3.10512

Quintero-Uribe, L. C., Navarro, L. M., Pereira, H. M., & Fernández, N. (2022). Participatory scenarios for restoring European landscapes show a plurality of nature values [Review]. Ecography, 2022(4), Article e06292. 10.1111/ecog.06292

Rawluk, A., Ford, R., Anderson, N., & Williams, K. (2019). Exploring multiple dimensions of values and valuing: a conceptual framework for mapping and translating values for social-ecological research and practice [Article]. Sustainability Science, 14(5), 1187–1200. 10.1007/s11625-018-0639-1

Raymond, C. M., Stedman, R., & Frantzeskaki, N. (2023). The role of nature-based solutions and senses of place in enabling just city transitions. Environmental Science & Policy, 144, 10–19. 10.1016/j.envsci.2023.02.021

Richardson, M., Dobson, J., Abson, D. J., Lumber, R., Hunt, A., Young, R., & Moorhouse, B. (2020). Applying the pathways to nature connectedness at a societal scale: a leverage points perspective. Ecosystems and People, 16(1), 387–401. 10.1080/26395916.2020.1844296

Robinson, M. A. (2018). Using multi-item psychometric scales for research and practice in human resource management. Human Resource Management, 57(3), 739–750.10.1002/hrm.21852

Runhaar, H., Runhaar, P., Bouwmans, M., Vink, S., Buijs, A., & Kleijn, D. (2019). The power of argument: Enhancing citizen’s valuation of and attitude towards agricultural biodiversity. International Journal of Agricultural Sustainability, 17(3), 231–242. 10.1080/14735903.2019.1619966

Schultz, P. W. (2001). The structure of environmental concern: Concern for self, other people, and the biosphere. Journal of Environmental Psychology, 21(4), 327-339. http://sfx.library.wur.nl:9003/sfx_local?sid=SP%3APSYI;id=pmid%3A;id=doi%3A10.1006%2Fjevp.2001.0227;issn=0272-4944;isbn=;volume=21;issue=4;spage=327;pages=327-339;date=2001;title=Journal%20of%20Environmental%20Psychology;atitle=The%20structure%20of%20environmental%20concern%3A%20Concern%20for%20self%2C%20other%20people%2C%20and%20the%20biosphere.;aulast=Schultz;pid=%3CAN%3E2002-10149-001%3C%2FAN%3E%3CAU%3ESchultz%2C%20P.%20Wesley%3C%2FAU%3E%3CDT%3Ejournal%3BPeer%20Reviewed%20Journal%3BEmpirical%20Study%3C%2FDT%3E

Schulz, C., & Martin-Ortega, J. (2018). Quantifying relational values — why not? Current Opinion in Environmental Sustainability, 35, 15–21. 10.1016/j.cosust.2018.10.015

See, S. C., Shaikh, S. F. E. A., Jaung, W., & Carrasco, L. R. (2020). Are relational values different in practice to instrumental values? Ecosystem Services, 44. 10.1016/j.ecoser.2020.101132

Spash, C. L. (2020). A tale of three paradigms: Realising the revolutionary potential of ecological economics. Ecological Economics, 169, 106518.

Stålhammar, S., & Thorén, H. (2019). Three perspectives on relational values of nature. Sustainability Science, 14(5), 1201–1212. 10.1007/s11625-019-00718-4

Steg, L. (2016). Values, Norms, and Intrinsic Motivation to Act Proenvironmentally. In Annual Review of Environment and Resources (Vol. 41, pp. 277–292).

Stern, P. C., Dietz, T., & Guagnano, G. Y. (1998). A brief inventory of values. Educational and psychological measurement, 58(6), 984–1001. http://sfx.library.wur.nl:9003/sfx_local?sid=Elsevier%3AScopus;genre=article;issn=00131644;volume=58;issue=6;spage=984;epage=1001;pages=984-1001;date=1998;title=Educational%20and%20Psychological%20Measurement;atitle=A%20brief%20inventory%20of%20values;aufirst=P.C.;auinit=P.C.;auinit1=P;aulast=Stern;artnum=;_service_type=getFullTxt

Taye, F. A., Vedel, S. E., & Jacobsen, J. B. (2018). Accounting for environmental attitude to explain variations in willingness to pay for forest ecosystem services using the new environmental paradigm. Journal of Environmental Economics and Policy, 7(4), 420–440. 10.1080/21606544.2018.1467346

Verschuuren, B., & Brown, S. (2019). Cultural and spiritual significance of nature in protected areas : governance, management and policy. Routledge. 10.4324/9781315108186

Wang, J., Zheng, R., Iabchoon, S., van Bodegom, P. M., Morpurgo, J., Remme, R. P., Hu, M., Tukker, A., Chen, W.-S., Huang, Y., Wang, Z., Li, C., & Cui, S. (2025). Social value-weighted greenspace exposure index: A novel metric integrating cultural ecosystem services for equitable benefits. Ecological Indicators, 180. 10.1016/j.ecolind.2025.114300

West, S., Haider, L. J., Masterson, V., Enqvist, J. P., Svedin, U., & Tengö, M. (2018). Stewardship, care and relational values [Review]. Current Opinion in Environmental Sustainability, 35, 30–38. 10.1016/j.cosust.2018.10.008

Williams, D. R., & Vaske, J. J. (2003). The Measurement of Place Attachment: Validity and Generalizability of a Psychometric Approach. Forest Science, 49(6), 830–840. http://www.scopus.com/scopus/inward/record.url?eid=2-s2.0-0347380937&partnerID=40&rel=R7.0.0

