## Supplementary material for "The “Plural Values of Nature” scale: An integrated psychometric scale to measure instrumental, intrinsic and relational values": Supplemetary material

**S1 Text: Methodological details on scale construction and data collection**

### Instrument development

An initial narrative literature review was carried out to identify existing related scales and explore relevant sub-dimensions that should be considered within each value dimension. Our focus in the scale development was on a clear theoretical and conceptual analysis of the three values, sensitive to potential overlaps and plurality. For instrumental values, we build on previous conceptualisations of ecosystem services (ESS) and nature's Contribution to People (NCP), differentiating between *material, non-material and regulating contributions* to people (Fig 2 **in full paper**)(Diaz et al., 2018). For intrinsic values, we also build on existing conceptualisations seeing *nature as an end in itself* and v*alue beyond human benefits* (Hourdequin, 2024). The conceptualisation of relational values proved to be the most difficult one, as this relatively new concept is still in development, is quite complex and broad, and has noticeable overlaps with non-instrumental anthropocentric values. Various scales and item-batteries exist to measure values of nature (e.g. Feucht et al., 2023; Klain et al., 2017; Lengieza et al., 2023; Mrotek et al., 2019; See et al., 2020). Meanwhile, to the best of our knowledge, no scale exists that meet all four criteria for a composite psychometric scale to measure the plural values of nature. Existing scales tend to focus on, or emphasize, only one value of nature, have not been explicitly validated, are developed and/or used with specific samples - most notably students, are focusing on nature as community, or they are developed and validated in only one country or one cultural context. We acknowledge that  several well established and cross-culturally validated scales exist which  attend to human-nature relationships, but not explicitly targeted at values of nature, such as the revised New Ecological Paradigm scale (NEP) (Dunlap et al., 2000); connectedness to nature scale (Mayer & Frantz, 2004), and the Environmental Concerns scale (Schultz, 2001).

Consequently, we decided to use these scales and items as inspiration and focused first on the identification of relevant subdimensions to relational values from literature, most notably the IPBES conceptualisation of relational values (Andersen et al., 2022). The IPBES Value Assessment differentiates between *Place attachment and identity*; *Fulfilment and well-being* (Eudemonia);

*Spirituality*; and *Care* (as stewardship responsibility) (Anderson, 2022). Based on other reviews of relational values (Feucht et al., 2023; Pratson et al., 2023), we added the notion of *Kinship* with non-human entities as a fifth subdimension. With the three subdimensions for instrumental values and two subdimensions for intrinsic values, this resulted in a structure to develop items consisting of in total ten sub-dimension (Fig 2 **in full paper**).

The initial formulations of items were created by two of the authors based on the narrative review of exiting items, and reformulating (and often simplifying) them to refer to each of the value dimensions. These formulations were validated in an online workshop with all co-authors to refine wording. During the workshop we evaluated the structure of the scale in three value dimensions consisting of ten sub-dimensions, whether the item formulations were appropriate operationalizations of these sub-dimensions, and whether the sub-dimensions were exhaustive and mutually exclusive. The question stem was "*When I think about nature in the future, I find it important that…*" and responses to each item had to be given on a 7-point scale from *"not important at all"* to *"very important".*

For a more conclusive semantic validation of the items, we carried out 16 cognitive interviews with people not professionally engaged in conservation or research (8 in the Netherlands and 8 in Denmark). Participants were requested to first give a response for all items, then to reflect on how understandable each item was, if they had any difficulties in answering, and how well they felt each item represented the associated value (i.e., instrumental, relational, intrinsic). Based on the findings, the wording of 15 items was simplified, two items were removed, and one was added (see S1 Table). All changes made to the instrument after this step are described in the result section and S1 Table. The pilot testing of the 28-items aimed at further refinement and shortening of the scale, resulting in 12 items in the main study and eventually 11 items in the final scale (Fig 3 **in full paper**).

### Data collections and samples

Scale validation was carried out with three independent data collections, each containing samples from multiple countries (Fig 1 **in full paper**). We aimed for sample sizes of at least 150 respondents to meet the recommended minimum sample size of five respondents per scale item to perform factor analysis (Hair, Black, et al., 2019).

For initial quantitative pre-testing the scale was translated from English to Dutch and Danish and checked by native speakers for semantic and conceptual equivalence. An online questionnaire was administered with a convenience sample of university students, some of these were international students, adding quality to the transnational scale development. During December 2023, 94 students were recruited from Wageningen University in the Netherlands and 101 students from the University of Copenhagen in Denmark. After further revisions of the scale (see results section, S1, S2, S3 Table) a pilot study was conducted February/March 2024 as an online questionnaire administered by a professional polling agency with a sample of the general public in the Netherlands (n=150), Spain (n=150) and Denmark (n=150) (S4 Table).

After final iterations of the scale, (see results section, S1, S5-S8 Table), including a back-translation of the survey to check for translation inconsistencies, it was administered October to November 2024 by a professional international polling agency as part of a larger online questionnaire on restoration and rewilding. The sample consisted of people from general public in the Netherlands (n=200), Spain (n=200), Sweden (n=200), Poland (n=200) and Romania (n=200) (S9 Table). The scale was translated into Dutch, Spanish, Swedish, Polish, and Romanian using the AI translation tool DeepL and checked by one or more native speakers for content and semantic validation, with quotas regarding gender, age, and education for each country, according to the most recent census information from the Eurostat database (European Commission, 2024) to achieve representative samples of residents aged 18 years or older. To answer some of the other research questions connected to the wider survey, an oversampling of rural residents in all countries was carried out by requesting 1/3 of respondents coming from predominantly rural regions, 1/3 from intermediate regions, and 1/3 from predominantly urban regions (see S9 Table). Ethical approval for the study was given by the Wageningen Research Ethics Committee for non-medical studies involving human subjects (WUR-REC-2024-112).

**S2 Text: Details of data analysis and validation**

For the statistical validation of the scale a range of common scale reliability and validity analysis (Hair, Gabriel, et al., 2019) were performed for each of the steps and samples (Fig 1 **in full paper**). Country samples were treated separately in the pre-testing and pilot study to support the cross-cultural adaptation and development of the scale (Beaton et al., 2000). All data analysis were carried out in *R*, and statistical significance was set at p=0.05.

<u>*Item analysis*</u>

Across all data collections we assessed the performance of individual items via basic descriptive statistics including mean values, standard deviations (SD), skewness, kurtosis, and response range to assess the clarity of items and response options and to check for potential ceiling or floor effects. Given that answers were recorded on a 7-point scale we considered mean values above 6.5 to indicate a ceiling effect and below 1.5 a floor effect. Regarding answer variability we defined low variability at SD <0.5 and high variability at SD > 2.0. If cases outside these levels were detected, we discussed if the results might be due to diverse perspectives on the issue or could be an effect of item wording that led to different interpretations by respondents. Regarding distributions, we inspected if the items were approximately normally distributed or showed strong deviations from normality considering values between -2 and +2 acceptable for skewness and kurtosis.

<u>*Exploratory factor analysis*</u>

Exploratory factor analysis (EFA) was used in the quantitative pre-testing and the pilot study to analyse the underlying factor structure and assess if the empirical results aligned with the theoretical model (i.e. intrinsic, instrumental and relational values). To verify the suitability of the data for EFA we inspected the Kaiser-Meyer-Olkin (KMO) Measure (> 0.60 indicates appropriateness of EFA) and if Bartlett's Test of Sphericity is significant (p < 0.05) indicating sufficient correlations among included variables (Hair, Gabriel, et al., 2019). We decided the number of retained factors based on eigenvalues >1 and chose an oblique rotation method (Direct Oblimin) as based on the theoretical framework we expected correlations between the underlying factors (i.e. values). We followed recommendations on statistical scale development by (Hair, Gabriel, et al., 2019) and excluded items that had communalities < 0.4 and KMO values < 0.5, indicating too low correlations with the other items. Several items showed moderate to strong deviations from normality, thus we chose to perform the EFA with Principal Axis Factoring (PAF), which is less sensitive to non-normality of data (Hair, Black, et al., 2019).

*Confirmatory factor analysis*

Confirmatory factor analysis was performed on the main survey to confirm the hypothesized three-factor model. Given that the main survey was comprised of samples from five countries we fitted a multi-Group CFA in *R* using the *lavaan* package (Rosseel, 2012). Multi-group CFA allows to test whether the factor structure is invariant across countries. Following common recommendations on multi-group CFA we initially tested the model on the entire dataset before adding country as a grouping variable. The appropriateness of our proposed models was assessed via multiple goodness-of-fit indices: we aimed for a Comparative Fit Index (CFI) and Tucker-Lewis Index (TLI) above 0.95, a Root Mean Square Error of Approximation (RMSEA) below 0.06 and a Standardized Root Mean Square Residual (SRMR) below 0.08, while standardized factor loadings should exceed 0.40 (Hair, Black, et al., 2019). CFA assumes multivariate normality of the data, based on the results of the item analysis this was not given; thus, we chose a robust version of the maximum likelihood estimator with scaled test statistics (equal to Yuan-Bentler) and robust standard errors (Huber-White) in our CFA (Rosseel, 2012). To assess measurement invariance across countries, we tested for configural invariance (same factor structure across countries), metric invariance (equal factor loadings across countries) and scalar invariance (equal item intercepts across countries).

*Reliability and validity*

The reliability of obtained factors was assessed via Cronbach's alpha, assuming values above 0.6 to indicate acceptable reliability (Hair, Gabriel, et al., 2019). Validity was assessed via the average variance extracted (AVE) which should be above 0.5 and inspection of standardized factor loadings, using the common threshold for acceptable factor loadings of 0.40. Discriminant validity analysis included the inspection of factor correlations and the Fornell-Larcker Criterion, which assesses if factors share more variance with their items (AVE value) than with any other factor (squared factor correlations) (Stensland et al., 2013). Given that we assumed the cooccurrence of multiple values within respondents (i.e. value plurality) we expected potential limitation in the discriminant validity of the scale

*Predictive validity of the final scale in the context of nature restoration*

Linear regression models were used to assess the predictive validity of the final scale. First, we created a composite scale for each of the three values by calculating the mean of all included items. The final study contained besides the *Plural Values of Nature* scale also several items measuring attitudes towards different restoration actions, asking how respondent would feel about these changes in their country and in an area close to them. We chose four attitude items ("Reducing the influence that humans have on nature in the country/area" and "Restoring natural processes and dynamics in the country/area"). The attitude items were then each used as dependent variable in a linear regression model, including the three composite scales (i.e. intrinsic, instrumental and relational value) as predictor variables. Predictive validity was assessed based on explained variance (adjusted $R^2$ value and the standardized regression coefficients β).

*Value typology*

To illustrate and better understand the plurality of values, we classified respondents on a typology inspired by the Nature-Futures Framework (Kim et al., 2023), consisting of all possible combinations of values. Classification was based on respondents' scores on the composite scales as these represent the priority given to the different values. Assuming values above 4.5 express a high priority, we classified respondents into the 8 combinatory typologies (Intrinsic; Instrumental; Relational; Intrinsic & Instrumental; Intrinsic & Relational; Instrumental & Relational; Intrinsic & Instrumental & Relational; None).

**S3 Text Extensive results pre-test and pilot study**

### Quantitative pre-testing

Data in the quantitative pre-test showed a high agreement with the importance of all presented items in both student samples (mean values > 4.4). Item analysis revealed a strong ceiling effect (mean scores above 6.5 on a 7-point scale) for three of the intrinsic value items and two of the instrumental value items representing regulating contributions (S2 Table). Given that most of the students were enrolled in nature conservation or management related programmes, we decided not to adjust the items based on this finding as mean values and distributions might be more moderate within more diverse samples.

For the Danish sample the initial KMO (0.779) and Bartlett's Test of Sphericity ($p<0.001$) showed the adequacy of performing and EFA. However, eight items had to be removed from the EFA due to too low item KMO and communalities (see S3 Table). After this, the EFA retained a 4-factor solution explaining 65% of the total variation within the dataset. The factor structure supports the theoretically expected three-dimensional structure of the scale with factor 1 consisting of the remaining 7 *relational value* items and factor 2 consisting of the 6 remaining *intrinsic value* items. However, *instrumental value* is split into two factors. The Dutch data was also suitable for an EFA (KMO=0.791, Bartlett's Test of Sphericity $p<0.001$), whereby six items were removed from the EFA due to too low communalities with the other items. Five factors were extracted which together explained 68% of the data variance. The factor structure was less clearly aligned with the assumed theoretical model than in the Danish pre-testing. However, sub-dimension grouped together with all items intended to represent *intrinsic value* and the *Care* sub-dimension building one factor, *spirituality* & *kinship* building the second factor, *material & regulating contributions* the third and *immaterial contributions*, *place attachment & identity* and *fulfilment & well-being* items the fourth.

Based on the findings of the statistical analysis and additional theoretical reasoning within the research team, a total of eight items were edited in order to improve their clarity and understandability for respondents (S1 Table).

### Pilot study

The pilot study returned representative samples of 150 complete responses from each of three included countries (S4 Table). Item analysis showed no strong ceiling effect (mean >6.5) for any of the items in either sample. However, the *Regulating contributions* items had high support and therefore low variability, with mean values above 6.0, less than 1-2% of responses on the negative end of the scale, and 80% of respondents choosing the two highest response options (S5 Table). Similar patterns were observed for the *Care* item and some of the intrinsic value items (S5 Table). Overall, the sample showed adequacy of performing an EFA and the initially retained solution for the complete dataset explained 62% of the variation with a four-factor structure (see S6 Table). Items representing intrinsic values build the first factor, confirming the assumed theoretical model regarding intrinsic values. However, the *Regulating contributions* items and the *Care* item and a relational item on local communities loaded also on the first factor (S6 Table). Taking the above-described findings into consideration while having the aim to significantly shorten the scale, we removed these items before performing an additional EFA for each country separately (S7 Table). The reduction in items resulted in the *Regulating contributions* sub-dimension within instrumental values and the *Care* sub-dimension within relational values no longer had any related items in the scale. The country based EFA also retained 4 factor solutions explaining 64%-69% of the variation (S7 Table). While the items that had to be removed due to too low communality varied across countries a general pattern showed that the items representing intrinsic values most clearly assembled in a common factor which had some cross loadings with the items from relational values in the Dutch and the Spanish sample (S7 Table). Some of the proposed factors also indicated an overlap between the instrumental and relational values. Based on the collective findings and analysis of the quantitative pretesting and pilot study we selected 13 of the pilot study items (see items in bold in S7 Table). Grouping them according to our theoretical model into instrumental, relational, and intrinsic value dimensions indicated good scale reliability (Cronbach alpha>0.70) across all countries (S8 Table). Given that the items representing *Material contributions* performed somewhat different across the countries (S7 Table) we decided to merge the item on harvesting wood with the item on collecting berries and mushrooms. This further shortened the scale to 12 items from eight sub-dimensions (S1 Table). In the hope of increasing the variability in answers we decided to change the questions stem from asking the respondent to indicate how important they felt various aspects of nature were when considering nature in the future, to asking them how much of a priority they felt the various aspects were when considering nature in the future (S1 Table).

**References**


Andersen, G., Fløttum, K., Carbou, G., & Gjesdal, A. M. (2022). People's Conceptions and Valuations of Nature in the Context of Climate Change [Article]. *Environmental Values*, *31*(4), 397-420. https://doi.org/10.3197/096327121X16328186623850

Anderson, C. B., Athayde, S., Raymond, C.M., Vatn, A., Arias, P., Gould, R.K., Kenter, J., Muraca, B., Sachdeva, S., Samakov, A., Zent, E., Lenzi, D., Murali, R., Balvanera, P., Pascual, U., Christie, M., Baptiste, B., and González-Jiménez, D. . (2022). Chapter 2: Conceptualizing the diverse values of nature and their contributions to people. In P. Balvanera, Pascual, U., Christie, M., Baptiste, B., and González-Jiménez, D. (Ed.), *Methodological Assessment Report on the Diverse Values and Valuation of Nature of the Intergovernmental Science-Policy Platform on Biodiversity and Ecosystem Services.* . IPBES secretariat. https://doi.org/10.5281/zenodo.6493134

Beaton, D. E., Bombardier, C., Guillemin, F., & Ferraz, M. B. (2000). Guidelines for the Process of Cross-Cultural Adaptation of Self-Report Measures. *Spine*, *25*(24). https://journals.lww.com/spinejournal/fulltext/2000/12150/guidelines_for_the_process_of_cross_cultural.14.aspx

Diaz, S., Pascual, U., Stenseke, M., Martin-Lopez, B., Watson, R. T., Molnar, Z., Hill, R., Chan, K. M. A., Baste, I. A., Brauman, K. A., Polasky, S., Church, A., Lonsdale, M., Larigauderie, A., Leadley, P. W., van Oudenhoven, A. P. E., van der Plaat, F., Schroter, M., Lavorel, S., . . . Shirayama, Y. (2018). Assessing nature's contributions to people. *Science*, *359*(6373), 270-272. https://doi.org/10.1126/science.aap8826

European Commission, E. (2024). https://ec.europa.eu/eurostat/web/main/home

Feucht, V., Dierkes, P. W., & Kleespies, M. W. (2023). The different values of nature: a comparison between university students' perceptions of nature's instrumental, intrinsic and relational values. *Sustainability Science*, *18*(5), 2391-2403. https://doi.org/10.1007/s11625-023-01371-8

Hair, J. F., Black, W. C., Babin, B. J., & Anderson, R. E. (2019). *Multivariate Data Analysis* (8th Edition ed.). Cengage Learning.

Hair, J. F., Gabriel, M. L. D. S., da Silva, D., & Braga Junior, S. (2019). Development and validation of attitudes measurement scales: fundamental and practical aspects. *RAUSP Management Journal*, *54*(4), 490-507. https://doi.org/10.1108/RAUSP-05-2019-0098

Hourdequin, M. (2024). *Environmental ethics: From theory to practice*. Bloomsbury Publishing.

Kim, H., Peterson, G. D., Cheung, W. W. L., Ferrier, S., Alkemade, R., Arneth, A., Kuiper, J. J., Okayasu, S., Pereira, L., Acosta, L. A., Chaplin-Kramer, R., den Belder, E., Eddy, T. D., Johnson, J. A., Karlsson-Vinkhuyzen, S., Kok, M. T. J., Leadley, P., Leclère, D., Lundquist, C. J., . . . Pereira, H. M. (2023). Towards a better future for biodiversity and people: Modelling Nature Futures. *Global Environmental Change*, *82*. https://doi.org/10.1016/j.gloenvcha.2023.102681

Klain, S. C., Olmsted, P., Chan, K. M. A., & Satterfield, T. (2017). Relational values resonate broadly and differently than intrinsic or instrumental values, or the New Ecological Paradigm. *PLoS ONE*, *12*(8), e0183962. https://doi.org/10.1371/journal.pone.0183962

Pratson, D. F., Adams, N., & Gould, R. K. (2023). Relational values of nature in empirical research: A systematic review. *People and Nature*, *5*(5), 1464-1479. https://doi.org/10.1002/pan3.10512

Rosseel, Y. (2012). lavaan: An R Package for Structural Equation Modeling. *Journal of Statistical Software*, *48*(2), 1-36. http://www.jstatsoft.org/v48/i02/

Stensland, S., Aas, O., & Mehmetoglu, M. (2013). The influence of norms and consequences on voluntary catch and release angling behavior [article]. *Human Dimensions of Wildlife*, *18*(5). https://doi.org/10.1080/10871209.2013.811617

**S1 Table. Overview of the item wording and the adaptations made during all steps of scale development and validation**. Italics indicates a change to the formulation from the previous step of validation. For the main study, the question stem was also modified.

| | **Step** | **Cognitive Interviews** | **Quantitative pre-testing** | **Pilot study** | **Main survey** |
|---|---|---|---|---|---|
| | **Question Stem** | *When I think about nature in the future, I find it important that…* | *When I think about nature in the future, I find it important that…* | *When I think about nature in the future, I find it important that…* | *Please indicate how much of a priority you feel each aspect should have when considering nature in your country in the future.* |
| **Value** | **Sub-dimension** | | | | |
| Instrumental | Material contributions | *it contributes to economic growth and gives job opportunities* | it contributes to economic growth and gives job opportunities | it contributes to economic growth and gives job opportunities | Nature contributes to economic growth and gives job opportunities |
| | | *opportunities to consume crops, wood and other materials are provided* | opportunities to harvest wood, berries, or mushrooms are provided | opportunities to collect berries, or mushrooms are provided | Nature offers opportunities to harvest goods such as wood, berries, mushrooms, or fish |
| | | | - | opportunities for harvesting wood are optimised | [Item removed] |
| | | *food and other materials are available to people* | [Item removed] | | |
| | Immaterial contributions | *outdoor recreation and sport opportunities are available* | outdoor recreation and sport opportunities are available | outdoor recreation and sport opportunities are available | Nature offers opportunities for outdoor recreation and sports |

| | | | | | |
|---|---|---|---|---|---|
| | | *physical and mental wellbeing is supported* | physical and mental wellbeing is supported | mental wellbeing is supported | Nature offers opportunities to enhance mental wellbeing |
| | | *the beauty of natural areas is preserved* | people can enjoy the beauty of natural areas | people can enjoy the beauty of natural areas | People can enjoy the beauty of natural areas |
| | | *peace and quietness can be found in nature* | people can find peace and quietness in nature | people can find peace and quietness in nature | [Item removed] |
| | Regulating contributions | *negative impacts of climate change, such as flooding, are reduced* | negative impacts of climate change, such as flooding, are reduced | negative impacts of climate change, such as flooding, are reduced | [Item removed] |
| | | *nature can help secure the quality of the air and the environment* | nature can help secure the quality of the air and the environment | nature can help secure the quality of the air | [Item removed] |
| | | | natural areas support agriculture, for example through pollinators | natural areas support agriculture, for example through pollinators such as bees and other insects | [Item removed] |
| Relational | Place attachment and identity | *a sense of belonging can be found in nature* | people can find a sense of belonging in nature | people can find a sense of belonging in nature | People can find a sense of belonging in nature |
| | | *it is recognised that nature is part of who we are as a community* | it is recognised that nature is part of local communities | it is recognised that nature is part of our local community | [Item removed] |
| | | *our heritage and history with nature are remembered* | the history of landscapes is acknowledged | the history of natural landscapes is acknowledged | [Item removed] |
| | Fulfilment and well-being | *nature contributes to getting fulfilment in life* | nature contributes to people's sense of fulfilment in life | nature contributes to people's sense of fulfilment in life | Nature contributes to people's sense of fulfilment in life |

| | | | | | |
|---|---|---|---|---|---|
| | | *I can find joy and satisfaction in contributing to positive developments to nature* | people can find joy and satisfaction in taking care of nature | people can find satisfaction in taking care of nature | [Item removed] |
| | Spirituality | *the religious, sacred and spiritual values of nature are respected* | the religious, sacred and spiritual values of nature are respected | the religious, sacred and spiritual values of nature are respected | The religious, sacred and spiritual values of nature are respected |
| | | *I can have a spiritual or religious connection with animals and trees* | people can have a religious or spiritual connection with animals and trees | people can have a religious or spiritual connection with animals and trees | [Item removed] |
| | Care | *we take care of nature out of a sense of responsibility* | we care for nature | people feel responsible for taking care of nature | [Item removed] |
| | | *we respect the characteristics and autonomy of all non-human living beings* | [Item removed] | | |
| | Kinship | *human beings are recognized as part of nature and nature as part of us* | *human beings are recognized as part of nature and nature as part of us* | human beings are recognized as part of nature and nature as part of us | [Item removed] |
| | | *I can relate to and feel connected with the trees and the animals* | *people can relate to and feel connected with the trees and the animals* | people can relate to and feel connected with the trees and the animals | People can relate to and feel connected with trees and animals |
| Intrinsic | Value beyond human benefits | *that the existence of nature without being harmed by people is respected* | *that the existence of nature without being harmed by people is respected* | the existence of nature without being harmed by people is respected | [Item removed] |
| | | *nature is protected regardless of whether this benefits humans or not* | *nature is protected regardless of whether this benefits humans or not* | nature is protected regardless of whether this benefits humans or not | Nature is protected regardless of whether this benefits humans or not |

| | | | | | |
|---|---|---|---|---|---|
| | | *individual non-human beings are protected regardless of whether this benefits humans or not* | *animals and plants are protected regardless of whether this benefits humans or not* | animals and plants are protected regardless of whether this benefits humans or not | [Item removed] |
| | | *our moral responsibility to protect nature is upheld* | *our moral responsibility to protect nature is upheld* | our moral responsibility to protect nature is upheld | Our moral responsibility to protect nature is upheld |
| | Nature as an end in itself | *nature is protected for its own sake* | *nature is protected for its own sake* | nature is protected for its own sake | Nature is protected for its own sake |
| | | *biodiversity is restored* | *biological diversity is restored* | biological diversity is restored | [Item removed] |
| | | *nature can flourish* | *nature is doing well* | nature is doing well | [Item removed] |
| | | *nature can develop freely and independent of human interest* | *nature can develop freely and independent of human interest* | nature can develop freely and independent of human interest | [Item removed] |

**S2 Table. Descriptive results from the quantitative pre-testing** with a convenience sample of university students. Bold indicates values with a ceiling effect (M>6.5)

| | | Sample | NL (n=94) | | | | | | DN (n=101) | | | | | |
|---|---|---|---|---|---|---|---|---|---|---|---|---|---|---|
| | | *When I think about nature in the future, I find it important that…* | M | SD | Min | Max | Sk | Ku | M | SD | Min | Max | Sk | Ku |
| Instrumental | 1 | *it contributes to economic growth and gives job opportunities* | 4.41 | 1.513 | 1 | 7 | -.495 | -.545 | 4.62 | 1.593 | 1 | 7 | -.495 | -.388 |
| | | *opportunities to harvest wood, berries, or mushrooms are provided* | 5.12 | 1.317 | 1 | 7 | -1.015 | 1.049 | 5.30 | 1.432 | 1 | 7 | -.882 | .080 |
| | 2 | *outdoor recreation and sport opportunities are available* | 6.04 | .978 | 2 | 7 | -1.412 | 3.178 | 6.00 | 1.279 | 1 | 7 | -1.684 | 3.087 |
| | | *physical and mental wellbeing is supported* | 5.60 | 1.100 | 1 | 7 | -1.220 | 2.871 | 5.65 | 1.236 | 1 | 7 | -1.316 | 2.812 |
| | | *people can enjoy the beauty of natural areas* | 5.89 | 1.065 | 2 | 7 | -1.259 | 2.169 | 5.94 | 1.066 | 2 | 7 | -1.092 | 1.321 |
| | | *people can find peace and quietness in nature* | 5.90 | 1.108 | 3 | 7 | -1.116 | .889 | 6.22 | 1.171 | 2 | 7 | -1.998 | 4.110 |
| | 3 | *negative impacts of climate change, such as flooding, are reduced* | 6.47 | .743 | 4 | 7 | -1.332 | 1.301 | **6.58** | **.791** | **2** | **7** | **-2.944** | **11.970** |
| | | *nature can help secure the quality of the air and the environment* | 6.33 | .687 | 5 | 7 | -.544 | -.765 | **6.70** | **.671** | **3** | **7** | **-3.005** | **11.083** |
| | | *natural areas support agriculture, for example through pollinators* | 5.46 | 1.138 | 2 | 7 | -.946 | .955 | 5.43 | 1.388 | 2 | 7 | -.873 | .111 |
| Relational | 4 | *people can find a sense of belonging in nature* | 5.30 | 1.243 | 2 | 7 | -.554 | -.166 | 5.56 | 1.212 | 2 | 7 | -.618 | -.140 |
| | | *it is recognised that nature is part of local communities* | 5.97 | .942 | 1 | 7 | -1.832 | 7.552 | 6.04 | 1.019 | 2 | 7 | -1.527 | 3.487 |
| | | *the history of landscapes is acknowledged* | 5.17 | 1.308 | 2 | 7 | -.569 | -.271 | 5.64 | 1.026 | 2 | 7 | -.596 | .771 |
| | 5 | *nature contributes to people's sense of fulfilment in life* | 5.18 | 1.503 | 1 | 7 | -.712 | .045 | 5.27 | 1.378 | 1 | 7 | -.800 | .335 |
| | | *people can find joy and satisfaction in taking care of nature* | 5.73 | 1.085 | 2 | 7 | -1.217 | 1.633 | 5.88 | 1.125 | 2 | 7 | -.922 | .540 |
| | 6 | *the religious, sacred and spiritual values of nature are respected* | 5.06 | 1.546 | 2 | 7 | -.468 | -.877 | 5.15 | 1.236 | 2 | 7 | -.224 | -.511 |
| | | *people can have a religious or spiritual connection with animals and trees* | 4.67 | 1.667 | 1 | 7 | -.548 | -.382 | 4.40 | 1.530 | 1 | 7 | -.235 | -.625 |
| | 7 | *we care for nature* | 6.41 | 1.003 | 1 | 7 | -2.951 | 11.855 | 6.45 | .880 | 2 | 7 | -2.200 | 6.513 |
| | 8 | *human beings are recognized as part of nature and nature as part of us* | 5.46 | 1.544 | 1 | 7 | -1.092 | .902 | 5.55 | 1.473 | 1 | 7 | -.952 | .525 |
| | | *people can relate to and feel connected with the trees and the animals* | 4.83 | 1.516 | 1 | 7 | -.621 | .064 | 4.89 | 1.503 | 2 | 7 | -.298 | -.669 |
| Intrinsic | 9 | *that the existence of nature without being harmed by people is respected* | 6.23 | 1.060 | 1 | 7 | -2.398 | 8.016 | 6.20 | 1.044 | 2 | 7 | -1.986 | 5.001 |
| | | *nature is protected regardless of whether this benefits humans or not* | 6.28 | 1.017 | 3 | 7 | -1.635 | 2.358 | 6.10 | 1.237 | 1 | 7 | -1.970 | 4.808 |
| | | *animals and plants are protected regardless of whether this benefits humans or not* | 6.29 | 1.114 | 2 | 7 | -1.844 | 3.030 | 6.28 | 1.087 | 1 | 7 | -2.289 | 6.985 |
| | | *our moral responsibility to protect nature is upheld* | **6.61** | **.676** | **4** | **7** | **-1.895** | **3.709** | **6.55** | **.866** | **2** | **7** | **-2.860** | **10.094** |
| | 10 | *nature is protected for its own sake* | 6.20 | .916 | 4 | 7 | -1.029 | .261 | 6.06 | 1.302 | 1 | 7 | -1.913 | 4.039 |
| | | *biological diversity is restored* | **6.71** | **.541** | **5** | **7** | **-1.753** | **2.221** | **6.58** | **.738** | **3** | **7** | **-2.348** | **6.841** |
| | | *nature is doing well* | **6.80** | **.547** | **4** | **7** | **-3.068** | **9.962** | **6.54** | **.944** | **1** | **7** | **-3.045** | **12.253** |
| | | *nature can develop freely and independent of human interest* | 6.11 | 1.126 | 2 | 7 | -1.768 | 3.435 | 5.82 | 1.291 | 2 | 7 | -1.252 | 1.296 |

**S3 Table. Results of the exploratory factor analysis from the quantitative pre-testing** with a convenience sample of university students. Values represent factor loadings after rotation, to ease reading small values (<.40) have been hidden.

| | | Sample | NL (n=94) | | | | | DN (n=90) | | | |
|---|---|---|---|---|---|---|---|---|---|---|---|
| | | **Factor** | **1** | **2** | **3** | **4** | **5** | **1** | **2** | **3** | **4** |
| | | *When I think about nature in the future, I find it important that…* | | | | | | | | | |
| Instrumental | 1 | *it contributes to economic growth and gives job opportunities* | | | .687 | | | | | .635 | |
| | | *opportunities to harvest wood, berries, or mushrooms are provided* | * | * | * | * | * | * | * | * | * |
| | 2 | *outdoor recreation and sport opportunities are available* | * | * | * | * | * | | | | .638 |
| | | *physical and mental wellbeing is supported* | | | | .589 | | | | .518 | |
| | | *people can enjoy the beauty of natural areas* | | | | .700 | | | | | .456 |
| | | *people can find peace and quietness in nature* | | | | .845 | | | | | .822 |
| | 3 | *negative impacts of climate change, such as flooding, are reduced* | | | .478 | | | * | * | * | * |
| | | *nature can help secure the quality of the air and the environment* | | | .517 | | | * | * | * | * |
| | | *natural areas support agriculture, for example through pollinators* | | | .478 | | | | | .728 | |
| Relational | 4 | *people can find a sense of belonging in nature* | | | | .629 | | .728 | | | |
| | | *it is recognised that nature is part of local communities* | * | * | * | * | * | * | * | * | * |
| | | *the history of landscapes is acknowledged* | | | | | .672 | * | * | * | * |
| | 5 | *nature contributes to people's sense of fulfilment in life* | | | | .560 | | .668 | | | |
| | | *people can find joy and satisfaction in taking care of nature* | | | | .431 | | .561 | | | |
| | 6 | *the religious, sacred and spiritual values of nature are respected* | .789 | | | | | .456 | | | |
| | | *people can have a religious or spiritual connection with animals and trees* | .862 | | | | | .828 | | | |
| | 7 | *we care for nature* | | -.723 | | | | * | * | * | * |
| | 8 | *human beings are recognized as part of nature and nature as part of us* | * | * | * | * | * | .747 | | | |
| | | *people can relate to and feel connected with the trees and the animals* | .539 | | | | | .794 | | | |
| Intrinsic | 9 | *that the existence of nature without being harmed by people is respected* | | -.816 | | | | * | * | * | * |
| | | *nature is protected regardless of whether this benefits humans or not* | | -.594 | | | | | .866 | | |
| | | *animals and plants are protected regardless of whether this benefits humans or not* | | -.670 | | | | | .786 | | |
| | | *our moral responsibility to protect nature is upheld* | * | * | * | * | * | | .606 | | |
| | 10 | *nature is protected for its own sake* | * | * | * | * | * | | .683 | | |
| | | *biological diversity is restored* | | -.640 | | | | | .668 | | |
| | | *nature is doing well* | | -.846 | | | | | .690 | | |
| | | *nature can develop freely and independent of human interest* | | -.624 | | | | * | * | * | * |
| *Eigenvalue* | | | 6.819 | 2.865 | 1.737 | 1.461 | 1.283 | 6.237 | 3.480 | 1.449 | 1.264 |
| ***Variance explained*** | | | **32.47** | **13.64** | **8.27** | **6.96** | **6.11** | **32.83** | **18.32** | **7.63** | **6.65** |
| *Correlation with factor 1* | | | 1 | -.263 | .166 | .396 | .148 | 1 | .323 | .310 | .476 |
| *Correlation with factor 2* | | | -.263 | 1 | -.154 | -.250 | -.161 | .323 | 1 | -.120 | .104 |
| *Correlation with factor 3* | | | .166 | -.154 | 1 | .219 | .051 | .310 | -.120 | 1 | .232 |
| *Correlation with factor 4* | | | .396 | -.250 | .219 | 1 | .148 | .467 | .104 | .232 | 1 |
| *Correlation with factor 5* | | | .148 | -.161 | .051 | .148 | 1 | - | - | - | - |

* removed because of KMO<0.5 or communality <0.4

**S4 Table. Sociodemographic characteristics of the pilot study samples**

| | Netherlands | | Spain | | Denmark | |
|---|---|---|---|---|---|---|
| | **Sample** | **CBS** | **Sample** | **CBS** | **Sample** | **EuroStat** |
| **Gender** | | | | | | |
| *Female* | 50% | 49% | 46% | 48% | 51% | 49% |
| *Male* | 50% | 51% | 54% | 52% | 49% | 51% |
| **Age** | | | | | | |
| *18-34* | 17% | 27% | 23% | 22% | 25% | 27% |
| *35-54* | 35% | 32% | 39% | 38% | 29% | 32% |
| *55-74* | 36% | 31% | 29% | 28% | 33% | 30% |
| *75+* | 13% | 10% | 9% | 12% | 13% | 11% |
| **Education** | | | | | | |
| *low* | 23% | 26% | 25% | 39% | 21% | 22% |
| *middle* | 41% | 42% | 35% | 25% | 42% | 42% |
| *high* | 35% | 32% | 40% | 36% | 36% | 36% |

***Note:***

We employed quota sampling, meaning screening criteria (i.e. gender, age, education, and region) were used to ensure respondents were a representative sample of residents of the Netherlands, Denmark and Spain aged 18 years or older. The Dutch sample also had a quota on urbanization. Data collection took place from February to March 2024 until target sample sizes and quotas were met.

**S5 Table. Descriptive results from the pilot study.** Bold indicates values with a ceiling effect (M>6.0)

| | | | Netherlands | | | | | | Denmark | | | | | | Spain | | | | | |
|---|---|---|---|---|---|---|---|---|---|---|---|---|---|---|---|---|---|---|---|---|
| | | | M | SD | MIN | MAX | SK | KU | M | SD | MIN | MAX | SK | KU | M | SD | MIN | MAX | SK | KU |
| Instrumental | 1 | it contributes to economic growth and gives job opportunities | 4.69 | 1.415 | 1 | 7 | -0.668 | 0.083 | 4.7 | 1.608 | 1 | 7 | -0.52 | -0.381 | 5.89 | 1.082 | 2 | 7 | -1.237 | 1.748 |
| | | opportunities to collect berries, or mushrooms are provided | 4.68 | 1.615 | 1 | 7 | -0.601 | -0.511 | 4.95 | 1.382 | 1 | 7 | -0.487 | 0.005 | 5.49 | 1.174 | 3 | 7 | -0.387 | -0.788 |
| | | opportunities for harvesting wood are optimised | 4.4 | 1.519 | 1 | 7 | -0.296 | -0.511 | 4.45 | 1.459 | 1 | 7 | -0.169 | -0.479 | 5.6 | 1.3 | 1 | 7 | -1.113 | 1.409 |
| | 2 | outdoor recreation and sport opportunities are available | 5.43 | 1.143 | 1 | 7 | -1.525 | 3.371 | 5.49 | 1.219 | 1 | 7 | -1.072 | 1.258 | 5.93 | 1.03 | 2 | 7 | -0.934 | 0.818 |
| | | mental wellbeing is supported | 5.41 | 1.281 | 1 | 7 | -1.378 | 2.266 | 5.47 | 1.191 | 1 | 7 | -0.866 | 1.231 | **6.2** | **1.056** | **1** | **7** | **-1.902** | **4.634** |
| | | people can enjoy the beauty of natural areas | 5.94 | 0.837 | 2 | 7 | -0.998 | 2.541 | 5.82 | 1.081 | 1 | 7 | -1.411 | 3.038 | **6.23** | **0.908** | **2** | **7** | **-1.462** | **2.867** |
| | | people can find peace and quietness in nature | 5.95 | 0.896 | 2 | 7 | -1.2 | 2.482 | 5.66 | 1.231 | 1 | 7 | -1.383 | 2.645 | **6.11** | **0.956** | **1** | **7** | **-1.709** | **5.41** |
| | 3 | negative impacts of climate change, such as flooding, are reduced | 5.9 | 1.145 | 1 | 7 | -1.594 | 3.742 | 5.91 | 1.17 | 2 | 7 | -1.181 | 1.117 | **6.19** | **1.149** | **1** | **7** | **-2.417** | **7.464** |
| | | nature can help secure the quality of the air | **6.2** | **0.777** | **4** | **7** | **-0.887** | 0.675 | **6.03** | **1.102** | **1** | **7** | **-1.623** | **3.629** | **6.4** | **0.969** | **1** | **7** | **-2.94** | **12.384** |
| | | natural areas support agriculture, for example through pollinators such as bees and other insects | **6.15** | **0.885** | **1** | **7** | **-1.822** | 7.017 | 5.77 | 1.1 | 2 | 7 | -0.826 | 0.42 | **6.29** | **0.854** | **3** | **7** | **-1.245** | **1.419** |
| Relational | 4 | people can find a sense of belonging in nature | 5.53 | 1.202 | 1 | 7 | -1.226 | 2.258 | 5.38 | 1.168 | 1 | 7 | -0.935 | 1.691 | 5.91 | 1.161 | 1 | 7 | -1.67 | 3.826 |
| | | it is recognised that nature is part of our local community | 5.86 | 0.934 | 2 | 7 | -0.917 | 1.194 | 5.77 | 1.094 | 2 | 7 | -0.847 | 0.486 | **6.07** | **1.072** | **1** | **7** | **-1.855** | **5.019** |
| | | the history of natural landscapes is acknowledged | 5.45 | 1.218 | 1 | 7 | -1.116 | 1.643 | 5.38 | 1.314 | 1 | 7 | -0.715 | 0.301 | **6.04** | **1.035** | **1** | **7** | **-1.551** | **3.691** |
| | 5 | nature contributes to people's sense of fulfilment in life | 5.32 | 1.368 | 1 | 7 | -1.379 | 2.271 | 5.44 | 1.184 | 2 | 7 | -0.863 | 0.777 | 5.79 | 1.168 | 1 | 7 | -1.472 | 3.178 |
| | | people can find satisfaction in taking care of nature | 5.15 | 1.387 | 1 | 7 | -0.924 | 0.601 | 5.77 | 1.044 | 3 | 7 | -0.825 | 0.44 | **6.14** | **0.941** | **1** | **7** | **-1.703** | **5.552** |

| | | | | | | | | | | | | | | | | | | | | |
|---|---|---|---|---|---|---|---|---|---|---|---|---|---|---|---|---|---|---|---|---|
| **Relational** | **6** | the religious, sacred and spiritual values of nature are respected | 4.13 | 1.797 | 1 | 7 | -0.294 | -0.858 | 4.63 | 1.548 | 1 | 7 | -0.51 | -0.131 | 4.95 | 1.575 | 1 | 7 | -0.809 | 0.156 |
| | | people can have a religious or spiritual connection with animals and trees | 3.68 | 1.837 | 1 | 7 | -0.084 | -1.064 | 4.21 | 1.574 | 1 | 7 | -0.338 | -0.375 | 4.89 | 1.767 | 1 | 7 | -0.647 | -0.349 |
| | **7** | people feel responsible for taking care of nature | **6.03** | **0.969** | **1** | **7** | **-1.579** | 4.5 | **6.06** | **0.985** | **3** | **7** | **-0.934** | **0.457** | **6.32** | **0.854** | **2** | **7** | **-1.783** | **4.99** |
| | **8** | human beings are recognized as part of nature and nature as part of us | 5.79 | 1.034 | 2 | 7 | -0.928 | 0.794 | 5.32 | 1.195 | 1 | 7 | -0.834 | 0.902 | 5.93 | 1.159 | 1 | 7 | -1.559 | 2.999 |
| | | people can relate to and feel connected with the trees and the animals | 5.05 | 1.441 | 1 | 7 | -0.844 | 0.532 | 4.92 | 1.531 | 1 | 7 | -0.695 | 0.106 | 5.81 | 1.276 | 1 | 7 | -1.469 | 2.354 |
| **Intrinsic** | **9** | the existence of nature without being harmed by people is respected | 5.93 | 1.047 | 1 | 7 | -1.75 | 5.433 | 5.88 | 1.22 | 1 | 7 | -1.362 | 2.367 | **6.26** | **1.006** | **1** | **7** | **-2.306** | **8.568** |
| | | nature is protected regardless of whether this benefits humans or not | 5.72 | 1.243 | 1 | 7 | -1.258 | 2.097 | 5.72 | 1.188 | 1 | 7 | -1.023 | 1.304 | 5.8 | 1.3 | 1 | 7 | -1.421 | 2.238 |
| | | animals and plants are protected regardless of whether this benefits humans or not | 5.47 | 1.335 | 1 | 7 | -1.144 | 1.713 | 5.85 | 1.126 | 1 | 7 | -1.137 | 1.865 | **6.1** | **1.219** | **1** | **7** | **-2.018** | **4.694** |
| | | our moral responsibility to protect nature is upheld | **6.08** | **0.98** | **1** | **7** | **-1.811** | **5.329** | 5.93 | 0.988 | 2 | 7 | -0.967 | 1.24 | **6.12** | **0.983** | **1** | **7** | **-1.793** | **5.677** |
| | **#** | nature is protected for its own sake | 5.79 | 1.095 | 1 | 7 | -1.601 | 4.251 | 5.77 | 1.292 | 2 | 7 | -1.049 | 0.658 | 5.82 | 1.248 | 1 | 7 | -1.521 | 2.842 |
| | | biological diversity is restored | 5.82 | 1.153 | 1 | 7 | -1.504 | 3.451 | 5.97 | 1.164 | 2 | 7 | -1.422 | 2.183 | **6.12** | **1.105** | **1** | **7** | **-2.239** | **6.946** |
| | | nature is doing well | 5.92 | 0.931 | 1 | 7 | -1.46 | 4.519 | **6.12** | **1.029** | **2** | **7** | **-1.217** | **1.422** | **6.37** | **0.931** | **1** | **7** | **-2.129** | **7.066** |
| | | nature can develop freely and independent of human interest | 5.63 | 1.144 | 1 | 7 | -1.07 | 1.589 | 5.58 | 1.119 | 2 | 7 | -0.742 | 0.764 | 5.85 | 1.252 | 1 | 7 | -1.412 | 2.273 |

**S6 Table. Initial result of the exploratory factor analysis from the pilot study.** Values represent factor loadings after rotation, to ease reading small values (<.40) have been hidden.

| | | | Total sample (n=450) | | | |
|---|---|---|---|---|---|---|
| | | Factor | 1 | 2 | 3 | 4 |
| Instrumental | | it contributes to economic growth and gives job opportunities | | 0.581 | | |
| | | opportunities to collect berries, or mushrooms are provided | | 0.416 | | |
| | 1 | opportunities for harvesting wood are optimised | | 0.78 | | |
| | | outdoor recreation and sport opportunities are available | | 0.446 | | |
| | | mental wellbeing is supported | | | | -0.417 |
| | | people can enjoy the beauty of natural areas | | | | -0.496 |
| | 2 | people can find peace and quietness in nature | | | | -0.697 |
| | | negative impacts of climate change, such as flooding, are reduced | 0.733 | | | |
| | | nature can help secure the quality of the air | 0.486 | | | |
| | 3 | natural areas support agriculture, for example through pollinators such as bees and other insects | * | * | * | * |
| Relational | | people can find a sense of belonging in nature | | | | -0.485 |
| | | it is recognised that nature is part of our local community | 0.513 | | | |
| | 4 | the history of natural landscapes is acknowledged | | | | |
| | | nature contributes to people's sense of fulfilment in life | | | | -0.42 |
| | 5 | people can find satisfaction in taking care of nature | | | | |
| | | the religious, sacred and spiritual values of nature are respected | | | -0.69 | |
| | 6 | people can have a religious or spiritual connection with animals and trees | | | -0.703 | |
| | 7 | people feel responsible for taking care of nature | 0.642 | | | |
| | | human beings are recognized as part of nature and nature as part of us | | | | -0.581 |
| | 8 | people can relate to and feel connected with the trees and the animals | | | -0.436 | |
| Intrinsic | | the existence of nature without being harmed by people is respected | 0.696 | | | |
| | | nature is protected regardless of whether this benefits humans or not | 0.87 | | | |
| | | animals and plants are protected regardless of whether this benefits humans or not | 0.872 | | | |
| | 9 | our moral responsibility to protect nature is upheld | 0.606 | | | |
| | | nature is protected for its own sake | 0.514 | | | |
| | | biological diversity is restored | 0.767 | | | |
| | | nature is doing well | 0.631 | | | |
| | 10 | nature can develop freely and independent of human interest | 0.637 | | | |
| Eigenvalue | | | 11.71 | 2.668 | 1.323 | 1.038 |
| **Variance explained** | | | 43.37 | 9.881 | 4.899 | 3.845 |
| Correlation with factor 1 | | | 1 | 0.289 | -0.258 | -0.622 |
| Correlation with factor 2 | | | 0.289 | 1 | -0.381 | -0.445 |
| Correlation with factor 3 | | | -0.26 | -0.38 | 1 | 0.243 |
| Correlation with factor 4 | | | -0.62 | -0.45 | 0.243 | 1 |

* removed because of communality <0.4

**S7 Table. Results of the exploratory factor analysis performed for each country within the pilot study.** Values represent factor loadings after rotation, to ease reading small values (<.40) have been hidden.

| | | | Netherlands (n=150) | | | | Denmark (n=150) | | | | Spain (n=150) | | | |
|---|---|---|---|---|---|---|---|---|---|---|---|---|---|---|
| | | **Factor** | **1** | **2** | **3** | **4** | **1** | **2** | **3** | **4** | **1** | **2** | **3** | **4** |
| **Instrumental** | | **it contributes to economic growth and gives job opportunities** | * | * | * | * | | 0.604 | | | 0.573 | | | |
| | | **opportunities to collect berries, or mushrooms are provided** | | 0.525 | | | | 0.408 | | | * | * | * | * |
| | 1 | **opportunities for harvesting wood are optimised** | | | | 0.571 | | 0.679 | | | * | * | * | * |
| | | **outdoor recreation and sport opportunities are available** | | | | 0.892 | * | * | * | * | 0.64 | | | |
| | | **mental wellbeing is supported** | 0.701 | | | | | | | | 0.546 | | | |
| | | **people can enjoy the beauty of natural areas** | 0.455 | | | 0.445 | | | -0.785 | | 0.542 | | | |
| | 2 | people can find peace and quietness in nature | * | * | * | * | | | -0.612 | | 0.888 | | | |
| **Relational** | | **people can find a sense of belonging in nature** | | | | | | | -0.649 | | 0.597 | | | |
| | 4 | the history of natural landscapes is acknowledged | 0.579 | | | | | | -0.42 | | | | -0.415 | |
| | | **nature contributes to people's sense of fulfilment in life** | 0.405 | 0.547 | | | | | -0.545 | | 0.521 | | | |
| | 5 | people can find satisfaction in taking care of nature | 0.415 | | | | | | -0.626 | | 0.556 | | | |
| | | the religious, sacred and spiritual values of nature are respected | | 0.858 | | | | | | -0.579 | | 0.903 | | |
| | 6 | **people can have a religious or spiritual connection with animals and trees** | | 0.88 | | | | | | -0.632 | | 0.79 | | |
| | | human beings are recognized as part of nature and nature as part of us | 0.565 | | | | * | * | * | * | 0.712 | | | |
| | 8 | **people can relate to and feel connected with the trees and the animals** | | 0.51 | | | | | | -0.471 | | | | |

| | | | | | | | | | | | | | | |
|---|---|---|---|---|---|---|---|---|---|---|---|---|---|---|
| Intrinsic | | the existence of nature without being harmed by people is respected | 0.444 | | | | 0.633 | | | | | | -0.772 | |
| | | nature is protected regardless of whether this benefits humans or not | | | -0.784 | | 0.82 | | | | | | -0.792 | |
| | | **animals and plants are protected regardless of whether this benefits humans or not** | | | -0.79 | | 0.868 | | | | | | -0.885 | |
| | 9 | **our moral responsibility to protect nature is upheld** | 0.639 | | | | 0.536 | | | | 0.427 | | | |
| | | **nature is protected for its own sake** | 0.403 | | -0.463 | | 0.643 | | | | | | | 0.709 |
| | | biological diversity is restored | 0.559 | | | | 0.667 | | | | | | -0.691 | |
| | | nature is doing well | 0.687 | | | | 0.435 | | | | | | -0.567 | |
| | 10 | nature can develop freely and independent of human interest | | | -0.752 | | 0.794 | | | | | | -0.558 | 0.462 |
| Eigenvalue | | | 9.226 | 2.449 | 1.398 | 1.214 | 11.71 | 2.668 | 1.323 | 1.038 | 10.635 | 1.587 | 1.169 | 1.109 |
| **Variance explained** | | | **43.92** | **11.66** | **6.655** | **5.78** | **43.371** | **9.881** | **4.899** | **3.845** | **50.641** | **7.555** | **5.564** | **5.282** |
| Correlation with factor 1 | | | 1 | 0.43 | -0.477 | 0.294 | 1 | 0.023 | -0.623 | -0.183 | 1 | 0.434 | -0.718 | 0.184 |
| Correlation with factor 2 | | | 0.43 | 1 | -0.282 | 0.418 | 0.023 | 1 | -0.301 | -0.359 | 0.434 | 1 | -0.384 | 0.254 |
| Correlation with factor 3 | | | -0.48 | -0.28 | 1 | -0.071 | -0.623 | -0.301 | 1 | 0.315 | -0.718 | -0.384 | 1 | -0.206 |
| Correlation with factor 4 | | | 0.294 | 0.418 | -0.071 | 1 | -0.183 | -0.359 | 0.315 | 1 | 0.184 | 0.254 | -0.206 | 1 |

**S8 Table. Reliability analysis of items selected for the final testing** (see Table S7). Cronbach alpha (ɑ) as measures of scale reliability.

| | Netherlands (n=150) | Denmark (n=150) | Spain (n=150) | Total (n=450) |
|---|---|---|---|---|
| **Instrumental** | 0.767 | 0.733 | 0.776 | 0.789 |
| **Relational** | 0.843 | 0.757 | 0.786 | 0.809 |
| **Intrinsic** | 0.809 | 0.791 | 0.707 | 0.765 |

**S9 Table. Sociodemographic characteristics of the main survey samples.**

| | Sweden | | Spain | | Romania | | Poland | | Netherlands | |
|---|---|---|---|---|---|---|---|---|---|---|
| | EuroStat | Sample | EuroStat | Sample | EuroStat | Sample | EuroStat | Sample | EuroStat | Sample |
| **Gender** | | | | | | | | | | |
| *Female* | 49.53% | 53.14% | 50.95% | 51.72% | 51.29% | 49.25% | 51.57% | 55.05% | 50.31% | 52.22% |
| *Male* | 50.47% | 45.89% | 49.05% | 48.28% | 48.71% | 50.75% | 48.43% | 44.95% | 49.69% | 47.29% |
| *Other* | NA | 0.97% | | | | | | | NA | 0.49% |
| **Age** | | | | | | | | | | |
| *18-34* | 20.49% | 23.19% | 17.65% | 20.2% | 18.22% | 20.6% | 18.38% | 23.23% | 20.33% | 21.18% |
| *35-49* | 18.25% | 23.67% | 21.58% | 29.56% | 21.31% | 27.64% | 22.78% | 26.77% | 17.07% | 22.17% |
| *50-64* | 17.51% | 24.15% | 20.91% | 27.09% | 19.83% | 28.14% | 18.02% | 24.75% | 19.67% | 26.11% |
| *65+* | 19.64% | 28.99% | 19.21% | 23.15% | 19.21% | 23.62% | 19.2% | 25.25% | 19.02% | 30.54% |
| **Rural-urban** | | | | | | | | | | |
| *intermediate* | 53.16% | 37.20% | 35.12% | 40.39% | 43.22% | 40.20% | 39.12% | 37.88% | 21.22% | 32.02% |
| *predominantly rural* | 6.94% | 28.50% | 2.66% | 24.63% | 44.76% | 30.65% | 38.10% | 30.81% | 0.59% | 22.66% |
| *predominantly urban* | 39.91% | 34.30% | 62.21% | 34.98% | 12.02% | 29.15% | 22.78% | 31.31% | 78.18% | 45.32% |

***Data weighing***

The final study used an oversampling of rural residents in all countries. To create more representative population estimates for the importance of the different values at the country level, the final dataset was weighted based on age, gender, and urban-rural residency for each country in accordance with Eurostat demographic data.

**S10 Table. Descriptive results from the main study**. Bold indicates values with a ceiling effect (M>6.5)

| | | ***Sample*** | **Sweden (N=208)** | | | | | | **Spain(N=204)** | | | | | | **Romania (N=207)** | | | | | | **Poland (N=201)** | | | | | | **Netherlands (N=208)** | | | | | |
|---|---|---|---|---|---|---|---|---|---|---|---|---|---|---|---|---|---|---|---|---|---|---|---|---|---|---|---|---|---|---|---|---|
| | | *When I think about nature in the future, I find it important that…* | **M** | **SD** | **Min** | **Max** | **Sk** | **Ku** | **M** | **SD** | **Min** | **Max** | **Sk** | **Ku** | **M** | **SD** | **Min** | **Max** | **Sk** | **Ku** | **M** | **SD** | **Min** | **Max** | **Sk** | **Ku** | **M** | **SD** | **Min** | **Max** | **Sk** | **Ku** |
| **Instrumental** | 1 | *Nature contributes to economic growth and gives job opportunities* | 5.38 | 1.371 | 1 | 7 | -0.675 | 0.089 | 5.60 | 1.370 | 1 | 7 | -1.078 | 1.303 | 5.71 | 1.482 | 1 | 7 | -1.154 | 0.875 | 5.55 | 1.264 | 1 | 7 | -0.741 | 0.536 | 4.64 | 1.386 | 1 | 7 | -0.495 | -0.055 |
| | | *Nature offers opportunities to harvest goods, such as wood, berries, mushrooms, or fish* | 6.08 | 1.120 | 1 | 7 | -1.496 | 2.746 | 5.81 | 1.277 | 1 | 7 | -0.994 | 0.818 | 6.27 | 1.062 | 1 | 7 | -2.021 | 5.441 | 5.86 | 1.235 | 1 | 7 | -1.043 | 0.784 | 4.99 | 1.438 | 1 | 7 | -0.712 | 0.282 |
| | 2 | *Nature offers opportunities for outdoor recreation and sports* | 5.88 | 1.177 | 2 | 7 | -0.886 | 0.084 | 6.06 | 1.152 | 1 | 7 | -1.679 | 3.534 | **6.56** | **0.760** | **4** | **7** | **-1.795** | **2.704** | 6.25 | 0.980 | 1 | 7 | -1.752 | 4.359 | 5.25 | 1.291 | 1 | 7 | -1.143 | 1.510 |
| | | *Nature offers opportunities to enhance mental wellbeing* | 6.09 | 1.037 | 2 | 7 | -1.226 | 1.602 | 6.17 | 1.079 | 1 | 7 | -1.525 | 2.717 | 6.43 | 0.877 | 3 | 7 | -1.641 | 2.368 | 6.22 | 0.992 | 1 | 7 | -1.610 | 3.748 | 5.53 | 1.199 | 1 | 7 | -0.993 | 1.211 |
| | | *People can enjoy the beauty of natural areas* | 6.07 | 1.086 | 2 | 7 | -1.288 | 1.765 | 6.18 | 1.097 | 1 | 7 | -1.451 | 2.176 | **6.50** | **0.897** | **1** | **7** | **-2.513** | **8.547** | 6.21 | 1.084 | 1 | 7 | -1.731 | 3.508 | 5.89 | 1.037 | 1 | 7 | -1.300 | 2.869 |
| **Relational** | 4 | *People can find a sense of belonging in nature* | 5.62 | 1.406 | 1 | 7 | -1.049 | 0.809 | 5.22 | 1.494 | 1 | 7 | -0.733 | 0.165 | 6.07 | 1.117 | 1 | 7 | -1.295 | 1.817 | 5.72 | 1.222 | 1 | 7 | -0.902 | 0.590 | 5.31 | 1.151 | 1 | 7 | -0.682 | 0.498 |
| | 5 | *Nature contributes to people's sense of fulfilment in life* | 5.82 | 1.275 | 1 | 7 | -0.990 | 0.623 | 5.43 | 1.432 | 1 | 7 | -0.873 | 0.558 | 6.17 | 1.136 | 1 | 7 | -1.511 | 2.319 | 5.63 | 1.227 | 1 | 7 | -0.699 | 0.512 | 5.28 | 1.308 | 1 | 7 | -1.009 | 1.283 |
| | 6 | *Nature provides religious, sacred and spiritual values or experiences* | 3.91 | 1.974 | 1 | 7 | 0.026 | -1.084 | 4.13 | 1.977 | 1 | 7 | -0.163 | -1.050 | 5.19 | 1.746 | 1 | 7 | -0.719 | -0.274 | 4.32 | 1.803 | 1 | 7 | -0.292 | -0.755 | 3.52 | 1.665 | 1 | 7 | -0.005 | -0.857 |
| | 8 | *People can relate to and feel connected with trees and animals* | 5.33 | 1.523 | 1 | 7 | -0.674 | -0.234 | 5.14 | 1.596 | 1 | 7 | -0.654 | -0.242 | 6.00 | 1.250 | 1 | 7 | -1.204 | 1.051 | 5.60 | 1.338 | 1 | 7 | -0.973 | 0.761 | 5.00 | 1.400 | 1 | 7 | -0.650 | 0.134 |
| **Intrinsic** | 9 | *Nature is protected regardless of whether this benefits humans or not* | 5.73 | 1.420 | 1 | 7 | -1.069 | 0.743 | 6.28 | 1.147 | 1 | 7 | -1.870 | 3.565 | 5.94 | 1.380 | 1 | 7 | -1.319 | 1.046 | 5.88 | 1.235 | 1 | 7 | -1.169 | 1.447 | 5.40 | 1.251 | 1 | 7 | -0.643 | 0.170 |
| | | *Our moral responsibility to protect nature is upheld* | 5.90 | 1.297 | 2 | 7 | -1.041 | 0.367 | 6.43 | 1.007 | 1 | 7 | -2.310 | 6.388 | 6.12 | 1.189 | 2 | 7 | -1.171 | 0.319 | 6.20 | 1.083 | 1 | 7 | -1.750 | 4.199 | 5.75 | 1.177 | 1 | 7 | -1.206 | 2.151 |
| | 10 | *Nature is protected for its own sake* | 5.82 | 1.299 | 1 | 7 | -1.086 | 0.967 | 6.35 | 1.027 | 1 | 7 | -1.951 | 4.388 | 6.16 | 1.273 | 1 | 7 | -1.786 | 3.124 | 5.94 | 1.240 | 1 | 7 | -1.350 | 2.271 | 5.72 | 1.248 | 1 | 7 | -1.045 | 1.384 |

**S11 Table. Multi-group CFA results for the 12 items.** Values represent standardized factor loadings for the hypothesized factor structure across all five samples, Cronbach alpha (ɑ), and Average Variance Extracted (AVE) as measures of scale reliability & validity

| **Values** | **Sweden (n=208)** | **Spain (n=204)** | **Romania (n=207)** | **Poland (n=201)** | **Netherlands (n=208)** |
|---|---|---|---|---|---|
| **Instrumental** | **ɑ = .854 AVE=.538** | **ɑ = .792 AVE=.428** | **ɑ = .706 AVE=.346** | **ɑ = .814 AVE=.439** | **ɑ = .769 AVE=.365** |
| *Nature contributes to economic growth and gives job opportunities* | 0.636 | 0.577 | 0.561 | 0.538 | 0.472 |
| *Nature offers opportunities to harvest goods, such as wood, berries, mushrooms, or fish* | 0.689 | 0.484 | 0.484 | 0.484 | 0.466 |
| *Nature offers opportunities for outdoor recreation and sports* | 0.767 | 0.702 | 0.645 | 0.764 | 0.49 |
| *Nature offers opportunities to enhance mental wellbeing* | 0.79 | 0.731 | 0.67 | 0.762 | 0.856 |
| *People can enjoy the beauty of natural areas* | 0.827 | 0.817 | 0.663 | 0.82 | 0.776 |
| ***Relational*** | **ɑ = .819 AVE=.511** | **ɑ = .770 AVE=.449** | **ɑ = .822 AVE=.544** | **ɑ = .769 AVE=.453** | **ɑ = .797 AVE=.478** |
| *People can find a sense of belonging in nature* | 0.835 | 0.71 | 0.813 | 0.853 | 0.815 |
| *Nature contributes to people's sense of fulfilment in life* | 0.848 | 0.829 | 0.815 | 0.829 | 0.808 |
| *Nature provides religious, sacred and spiritual values or experiences* | 0.559 | 0.528 | 0.656 | 0.42 | 0.52 |
| *People can relate to and feel connected with trees and animals* | 0.734 | 0.687 | 0.755 | 0.727 | 0.703 |
| ***Intrinsic*** | **ɑ = .841 AVE=.642** | **ɑ = .859 AVE=.668** | **ɑ = .750 AVE=.526** | **ɑ = .774 AVE=.536** | **ɑ = .820 AVE=.602** |
| *Nature is protected regardless of whether this benefits humans or not* | 0.827 | 0.78 | 0.786 | 0.699 | 0.781 |
| *Our moral responsibility to protect nature is upheld* | 0.805 | 0.816 | 0.75 | 0.739 | 0.784 |
| *Nature is protected for its own sake* | 0.765 | 0.864 | 0.622 | 0.759 | 0.764 |
| **Instrumental x Relational** | 0.931 | 0.798 | 0.899 | 0.885 | 0.929 |
| **Instrumental x Intrinsic** | 0.674 | 0.851 | 0.769 | 0.892 | 0.658 |
| **Relational x Intrinsic** | 0.689 | 0.64 | 0.757 | 0.787 | 0.719 |

**S12 Table. Value typology distribution** across the five countries and the total sample

| Typology | Sweden | Spain | Romania | Poland | Netherlands | TOTAL |
|---|---|---|---|---|---|---|
| *Intrinsic* | 6.70% | 4.90% | 0.50% | 3.00% | 10.30% | 5.10% |
| *Instrumental* | 8.20% | | 4.50% | 2.50% | 3.90% | 3.90% |
| *Relational* | 0.50% | | | | | 0.10% |
| *Intrinsic & Instrumental* | 19.20% | 30.90% | 8.00% | 17.30% | 22.10% | 19.60% |
| *Intrinsic & Relational* | 1.40% | 1.00% | 2.00% | 1.50% | 8.30% | 2.90% |
| *Instrumental & Relational* | 3.40% | | 1.00% | 0.50% | 0.50% | 1.10% |
| *Intrinsic & Instrumental & Relational* | 57.70% | 57.80% | 82.40% | 71.10% | 49.50% | 63.50% |
| *NEITHER* | 2.90% | 5.40% | 1.50% | 4.10% | 5.40% | 3.90% |
